# Transcriptional reprogramming in fused cells is triggered by plasma-membrane diminution

**DOI:** 10.1101/832378

**Authors:** Daniel Feliciano, Isabel Espinosa-Medina, Aubrey Weigel, Kristin M. Milano, Zhonghua Tang, Tzumin Lee, Harvey J. Kliman, Seth M. Guller, Carolyn M. Ott, Jennifer Lippincott-Schwartz

## Abstract

Developing cells divide and differentiate, and in many tissues, such as bone, muscle, and placenta, cells fuse acquiring specialized functions. While it is known that fused-cells are differentiated, it is unclear what mechanisms trigger the programmatic-change, and whether cell-fusion alone drives differentiation. To address this, we employed a fusogen-mediated cell-fusion system involving undifferentiated cells in tissue culture. RNA-seq analysis revealed cell-fusion initiates a dramatic transcriptional change towards differentiation. Dissecting the mechanisms causing this reprogramming, we observed that after cell-fusion plasma-membrane surface area decreases through increased endocytosis. Consequently, glucose-transporters are internalized, and cytoplasmic-glucose and ATP transiently decrease. This low-energetic state activates AMPK, which inhibits YAP1, causing cell-cycle arrest. Impairing either endocytosis or AMPK prevents YAP1 inhibition and cell-cycle arrest after fusion. Together these data suggest that cell-fusion-induced differentiation does not need to rely on extrinsic-cues; rather the plasma-membrane diminishment forced by the geometric-transformations of cell-fusion cause transient cell-starvation that induces differentiation.

## Introduction

Rapidly induced changes in fundamental cellular features such as cell shape, subcellular organization and cell bioenergetics can drive adaptive regulatory mechanisms ensuring cell survival. These changes also impact whether a cell undergoes cell proliferation or acquires a specialized function (Basson, 2012; McBeath et al., 2004; Paluch and Heisenberg, 2009; Tatapudy et al., 2017). Cell fusion, an essential process during regeneration and development, is a remarkable example of how cellular morphogenesis arising from the merging of two or more cells influences cell fate determination (Ogle et al., 2005; Oren-Suissa and Podbilewicz, 2007; Zhou and Platt, 2011), but whether the cellular changes in response to cell fusion are sufficient to promote a transcriptional program which supports the new differentiated state remains unknown.

In mammals, cell fusion occurs in different tissues and organs including bone, skeletal muscle, immune cells, and placenta (Oren-Suissa and Podbilewicz, 2007; Zhou and Platt, 2011). These systems have evolved specific fusion proteins, or fusogens, that allow precise modulation of membrane fusion events. Upon fusion, plasma membranes and cytoplasmic content from fusing cells are combined to form the new syncytium. The resulting multi-nucleated syncytia can be comprised of hundreds to millions of cells, creating a unique cellular environment in which individual nuclei stop dividing and differentiation programs are deployed to promote overall homeostasis within the fused cells (Mayhew and Simpson, 1994)(Goldman-Wohl and Yagel, 2014; Zhou and Platt, 2011). This physical transformation suggests an underlying process controlling syncytial differentiation. In *in vivo* settings, specification of each syncytium occurs in a complex environment containing extracellular signaling molecules. This has led to the view that syncytial differentiation is solely dependent on such extrinsic factors. However, it is possible that the unification of multiple cells by itself invokes cell-intrinsic pathways that contribute to their transcriptional reprogramming and differentiation.

Prior work has shown that cell fusion results in significant alterations in fundamental cell biological characteristics. These include changes in surface expression of membrane proteins, organelle intermixing, repositioning of nuclei, hormone secretion, and variations in metabolism including the regulation of the AMP-activated protein kinase (AMPK) (Bax and Bloxam, 1997; Cadot et al., 2015; Calvert et al., 2016; Costa, 2016; Coutifaris et al., 1991; Feliciano et al., 2018; Ferris et al., 1987; Finley, 2018; Niesler et al., 2007; Pellett et al., 2013; Riquelme, 2011; Ryall, 2013; Towler et al., 2004; Villee, 1969; Wang et al., 2014). It is unclear whether these and/or other changes function as cell-intrinsic cues required for subsequent downstream differentiation after cell fusion. This knowledge gap stems from the challenge of dissecting the distinctive contribution of cell fusion from differentiation signals within tissues.

To circumvent these constraints, we employed a VSV-G fusogen-mediated assay to rapidly induce cell fusion of undifferentiated culture cells in the absence of tissue-specific differentiation cues. Using this approach, we studied the structural, subcellular and transcriptional changes modifying the resulting syncytium, and explored the molecular mechanisms involved. We showed that VSV-G mediated cell fusion replicates hallmarks of *in vivo* fusing systems. Surprisingly, we found that the act of cell fusion alone induces changes in gene expression, inhibiting cell proliferation while favoring a differentiated cell state. This transcriptional reprogramming is dependent on remodeling of the plasma membrane landscape and an acute low energy state leading to the activation of AMPK and the downstream inhibition of YAP1.

## Results

### Cell fusion in a model system recapitulates physiological syncytial hallmarks

To determine whether the initial changes induced by cell fusion contribute directly to the fate of a syncytium, cellular alterations solely induced by cell fusion need to be isolated from differentiation cues existing within tissues. To accomplish this, we employed an *in vitro* **V**esicular **S**tomatitis **V**irus **G** protein (VSV-G) mediated fusion assay to trigger cell fusion in culture cells (Feliciano et al., 2018; Gottesman et al., 2010; Pellett et al., 2013). In this assay, VSV-G expressing cells were rapidly washed (5-10 seconds) with an isotonic, low pH buffer (Fusion Buffer), which activated VSV-G to induce fusion of plasma membranes of two or more adjacent cells within seconds (Supplementary Figure 1A). Visualizing different fluorescent subcellular markers, we then measured the degree that cell fusion re-shaped fundamental cellular features such as plasma membrane (PM), cytoplasmic organization, and the cell-cycle state.

To assess how quickly and efficiently cell fusion occurred upon induction, we examined the speed of exchange of cytoplasmic proteins between fusing cells. For this we monitored fluorescence intensity changes after fusing SUM-159 cells expressing only VSV-G (Receiver cells) or both VSV-G and a fluorescent cytoplasmic marker (Donor cells) (Figure 1A, B, Supplementary Figure 1B and Supplementary Video 1). Within 30-60 seconds after cell fusion initiation, the fluorescence intensity of Receiver cells increased while Donor cell intensities started to decrease. This reflected the formation of fusion pores and the beginning of cytoplasmic mixing (indicated as Fusion in Figure 1B). Full equilibration of the fluorescent cytoplasmic marker across the synctium was achieved after 7-10 minutes (Figure 1A, B, Supplementary Figure 1B). The exchange of large subcellular compartments took longer (Supplementary Figure 1C-D).

**Figure 1.**
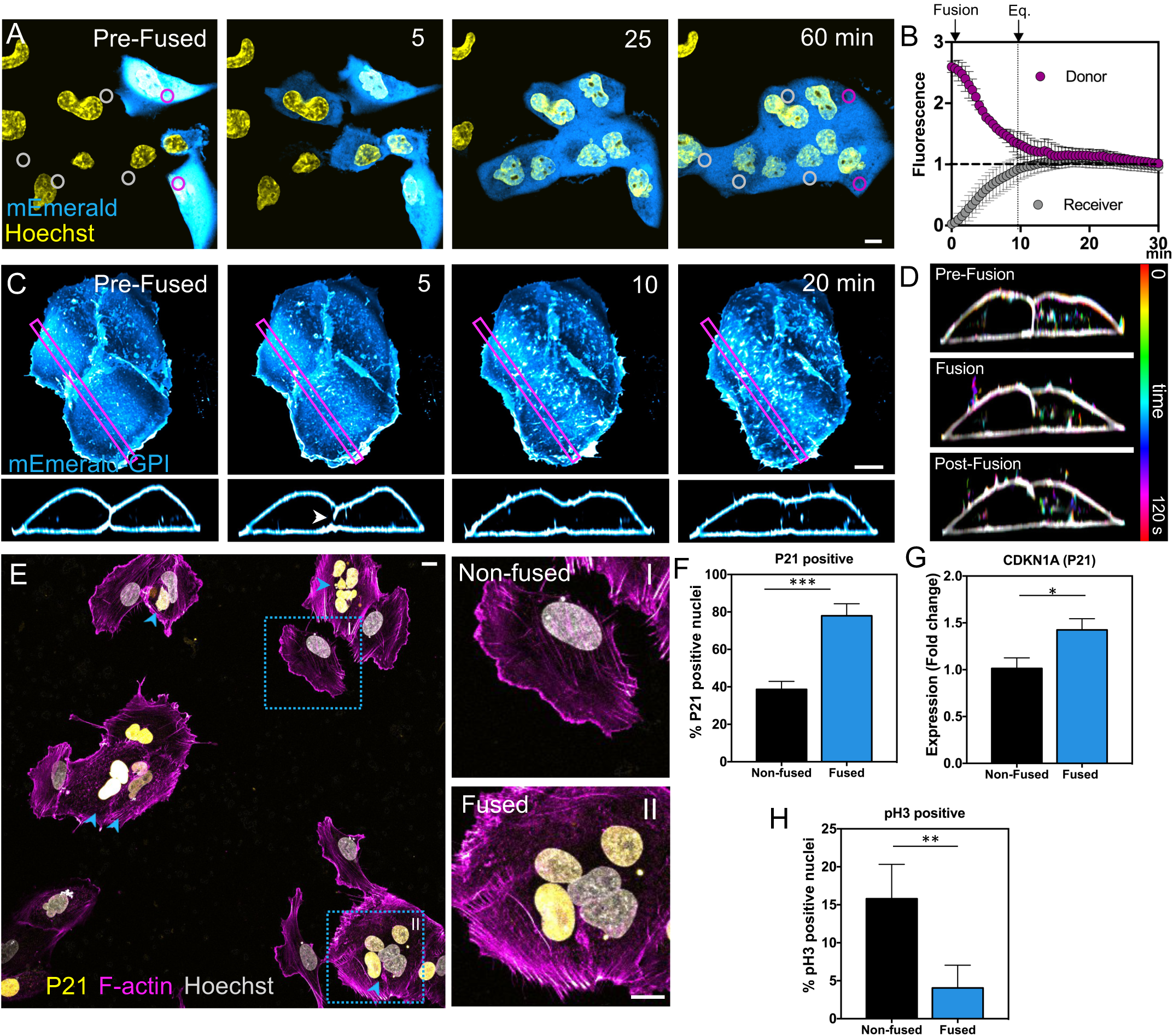
Changes in fundamental cellular features upon cell fusion lead to cell cycle exit. (A-B) Cells expressing VSV-G alone (Receiver cells) or co-expressing VSV-G and a cytoplasmic marker (Donor cells, cytoplasmic mEmerald) were mixed and fused by a brief wash with Fusion Buffer and then were imaged by confocal microscopy. Cytoplasmic mixing was measured as fluorescence intensity changes in ROIs of Donor (magenta ROI) and Receiver (gray ROI) cells over-time upon fusion (See also Supplementary Video 1). (C and D) Cells stably expressing VSV-G and a plasma membrane marker (mEmerald-GPI) were induced to fuse. Fusion-pore formation (C, insets, white arrow head) and changes in the plasma membrane were assessed by lattice-light sheet microscopy. (D) Temporal color-coded images were generated by compressing 2 minutes from time points before (Pre-fusion), during (Fusion), and after (Post-fusion) cell fusion (See also Supplementary Video 2). (E) Representative images of immunofluorescence staining of P21 positive nuclei in non-fused and fused cells (insets I and II, respectively) (Arrow heads point to positive P21 nuclei in fused cells). (F) The percentage of P21 positive nuclei in non-fused and fused cells were quantified 24hr after washing with Fusion Buffer. (G) Fold change in CDKN1A transcript (which codes for P21) expression was quantified after qRT-PCR in non-fused and fused cells. (H) Non-fused and fused cells were stained for the protein pH3, a positive indicator of mitosis, and the percentage of pH3 positive nuclei was quantified 24hr after washing with Fusion Buffer. Means ± SEM are shown, (***) P < 0.0001, (**) P = 0.03, (*) P < 0.05, Scale bar size = 10μm

Plasma membrane remodeling at the interface of fusing cells was examined using lattice light sheet microscopy (Figure 1C, D). Cells stably expressing the fluorescent plasma membrane marker, glycosylphosphatidyl-inositol anchored enhanced green fluorescent protein (GPI-EGFP), and transfected with VSV-G were rapidly washed with Fusion Buffer to induce fusion of plasma membranes of two or more adjacent cells. Within seconds the plasma membranes began to rearrange (Figure 1C, D). Over several minutes the membrane boundary between adjacent cells disappeared as their plasma membranes coalesced (Figure 1C lower panel, and Supplementary Video 2). During this process, the boundary between the two cells disappeared first in a small area near the bottom of the cell and then propagated upward until only one cell outline was visible. We also observed other morphological changes in the plasma membrane including dynamic membrane ruffling and the appearance of membrane projections emerging approximately 3-5 min after triggering cell fusion (Figure 1C, D, Supplementary Video 2). Correlating with the dramatic changes in PM remodeling were changes in actin dynamics (Supplementary Video 4).

Nuclei tracking analyses further demonstrated that nuclei from fusing cells quickly start to congregate at the center of the newly formed syncytium after 10 minutes, forming a less-dynamic cluster of nuclei 60 min after cell fusion (Supplementary Figure 1E, F, Supplementary Video 5). Importantly, several of these subcellular changes, such as cytoplasmic mixing and nuclear clustering, have also been described *in vivo* after macrophage fusion and during the development of skeletal muscle and placenta (Cadot et al., 2015; Calvert et al., 2016; Jeganathan et al., 2014; Wang et al., 2014).

Another representative feature shared by different syncytia is the loss of their competence to enter the cell cycle (Chuprin et al., 2013; Duelli and Lazebnik, 2003; Goldman-Wohl and Yagel, 2014). To test whether VSV-G mediated cell fusion leads to cell cycle arrest, we measured the levels of the cell cycle arrest marker P21 (Sherr and Roberts, 1995). We found that fused cells experience a 2-fold increase in P21 positive nuclei 24 hr after induction of cell fusion (Figure 1E, F). In addition, expression of P21 transcripts (CDKN1A), measured by qRT-PCR, increases in fused cells (Figure 1G). In contrast, the nuclear levels of the positive mitotic marker pH3 are reduced by 3-fold in fused cells (Figure 1H). In agreement with these results, after monitoring newly formed syncytia for 36 hr, we confirmed that 95% of fused cells did not undergo cell division (Supplementary Figure 2). These findings are consistent with an arrest in mitotic entry in fused cells and suggest that the act of cell fusion, alone, can initiate transcriptional changes.

Altogether, these observations show in detail how cell fusion quickly remodels fundamental cellular features including cell shape (through PM and cytoskeleton dynamics), subcellular organization (through cytoplasmic and organelle intermixing), and the cell-cycle state of a newly formed syncytium (directly restricting its capacity to enter the cell-cycle). Furthermore, these results demonstrate that VSV-G mediated cell fusion is a suitable model to test whether the changes induced by this process trigger cell-intrinsic mechanisms modulating syncytial function, as the characteristics observed in this model system recapitulated those seen in physiological syncytial systems.

### Cell fusion is sufficient to induce transcriptional reprogramming towards a differentiated state

Cell-cycle arrest is a representative characteristic that is coordinated with the reprogramming of gene expression during terminal differentiation (Buttitta and Edgar, 2007). To test whether cell fusion is sufficient to induce transcriptional reprogramming we performed RNA-SEQ of VSV-G fused and non-fused cells (Figure 2A). Differential expression analyses using False discovery rate (FDR)<0.05, revealed 2,513 genes that differed between fused and non-fused cells. Functional annotation clustering using DAVID Bioinformatics Resources (Huang da et al., 2009) revealed that the majority of these genes are enriched in clusters of plasma membrane, vesicular, and cytoplasmic cell component genes (Figure 2B), suggesting an overall structural remodeling that supports a new cellular state in fused cells.

**Figure 2.**
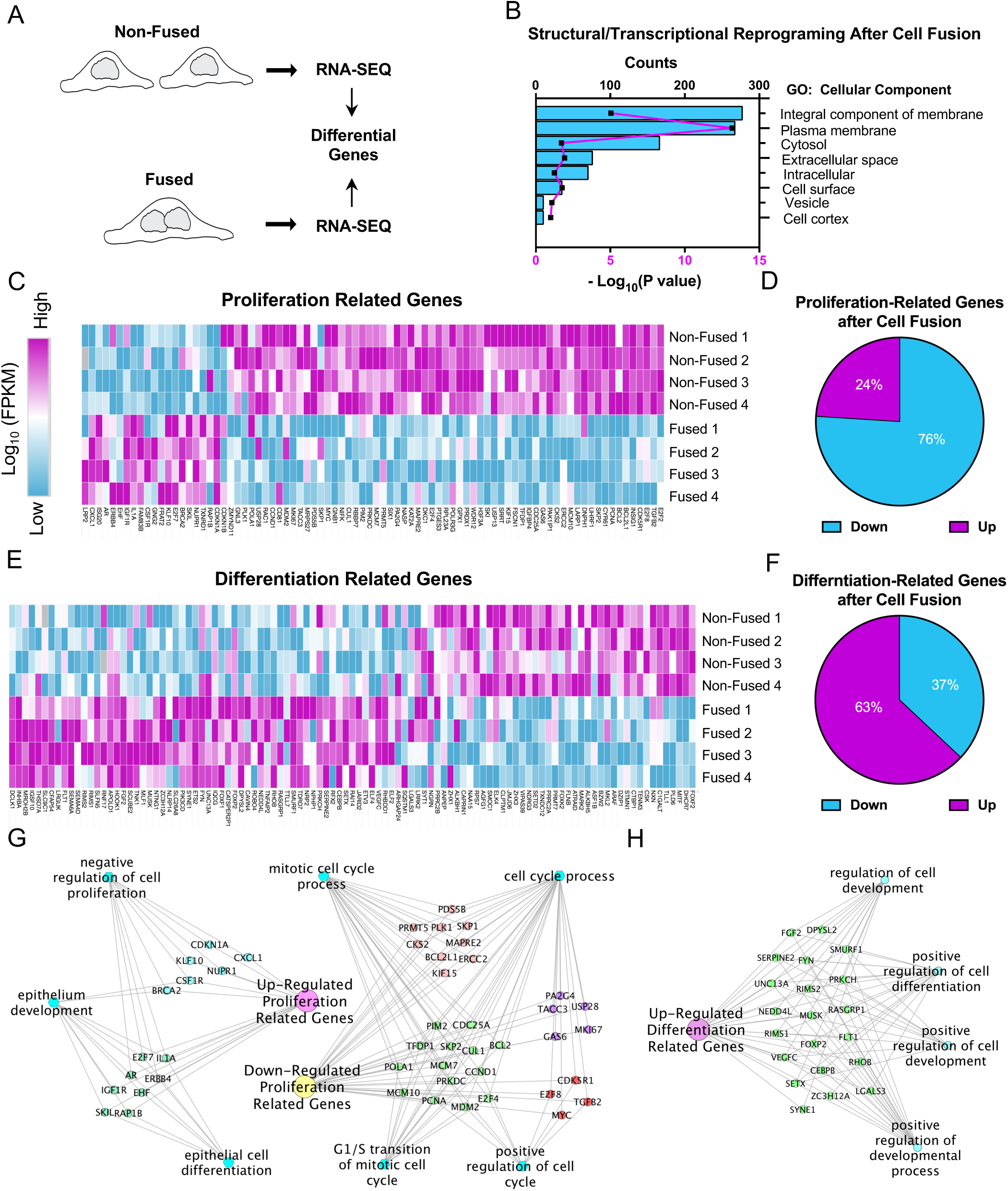
Cell fusion decreases the expression of proliferation-related genes while promoting expression of genes involved in differentiation and cell-cycle arrest. (A) Graphical description of RNAseq work flow to compare fused and non-fused SUM159 cell transcriptional profiles 6hr after induction of cell fusion. (B) Genes differentially expressed between fused and non-fused cells were filtered by gene ontologies (GO). The differentially expressed genes were grouped into Cellular Component genes that significantly changed after cell fusion. (C - F) Differentially expressed genes were filtered for both cell proliferation-related (C)(GO 0008283) and cell differentiation-related (E)(GO 0030154) transcripts. Expression levels (Log10 FPKM) of 4 independent experiments are shown as heatmap visualizations. Comparison of the percentage of proliferation (D) and differentiation (F) genes that were down or up-regulated after cell fusion. (G and H) ToppCluster plots showing the functional network among proliferation and differentiation related differentially expressed genes.

To assess if fused cells were changing their gene expression profile towards a differentiated state, we searched among all differentially expressed genes for either cell proliferation (GO: 0008283) or cell differentiation (GO: 0030154) related genes (Figure 2C-F). Hierarchical clustering of these genes showed that a large cluster comprising 76% of total cell proliferation genes was down-regulated (Figure 2C, D). Conversely, 63% of cell differentiation genes were up-regulated in fused cells (Figure 2E, F). Global gene regulatory network analysis showed that up-regulated cell differentiation genes are sub-grouped into genes that promote both differentiation and development (Figure 2H). Furthermore, in addition to down-regulated genes promoting the cell cycle, a subgroup of proliferation related genes was up-regulated. These genes are classified as negative regulators of cell proliferation, including P21 (CDKN1A) (Figure 2G). This is consistent with our results of syncytial cell cycle arrest (Figure 1E-H) and demonstrates that cell fusion is sufficient to induce transcriptional reprogramming favoring a cell differentiated state.

### Cell fusion blocks the expression of proliferation genes by YAP1 redistribution from the nucleus into cytoplasm

While negative cell-cycle regulators promote differentiation, positive cell cycle inducers prevent it. One of these inducers is the Yes-associated-protein-1 (YAP1), which when active promotes cell proliferation (Zhu et al., 2015b). Interestingly, specific YAP1 downstream target genes were also part of the list of differentially expressed genes identified by our RNA-SEQ analyses (Figure 3A). Consistent with cell-cycle arrest and the acquisition of a differentiated state, most of these genes where down-regulated by 15-45% in fused cells, suggesting YAP1 activity might be negatively regulated upon cell fusion (Figure 3B).

**Figure 3.**
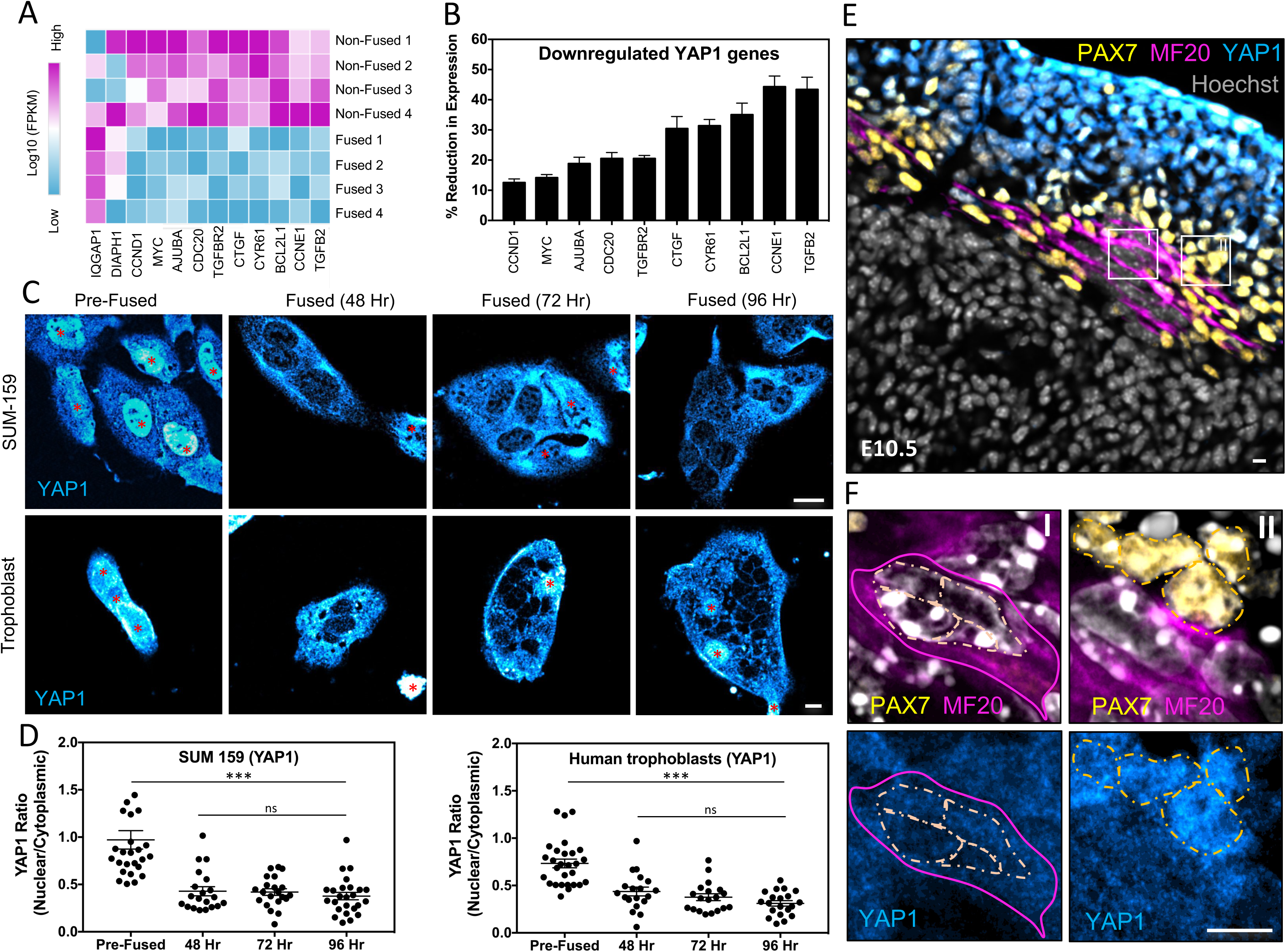
Cell fusion leads to YAP1 inhibition and cytoplasmic localization. (A) Heatmap representing the expression levels (Log10 FPKM) of transcripts downstream of YAP1 in non-fused and fused cells (4 independent experiments). (B) Average reduction of indicated transcripts downstream of YAP1 in fused cells. (C and D) Immunofluorescence was used to determine the localization of YAP1 in VSV-G transfected SUM-159 cells, either before or after fusion (C, top panel); and in isolated human trophoblast cells that fuse in culture without induction (D, bottom panel). Representative images are shown. Red * marks non-fused cells. (D) The nuclear/cytoplasmic ratio of YAP1 before and after (48, 72, and 96 hr) cell fusion in SUM-159 cells and human trophoblasts was calculated. (E and F) 25 μm sections of mouse E10.5 skeletal muscle were fixed and immunostained to determine YAP1 localization. Progenitor muscle cells (non-fused) were labeled using PAX7 antibodies while fused muscle cells were labeled using MF20 antibodies. Light-orange outlines surround the nuclei in fused cells (F, inset I) or progenitor cells (F, inset II). A magenta outline illustrates the boundary of a fused muscle cell (inset I). Means ± SEM are shown, (***) P < 0.0001, (no significant = ns) P > 0.05, Scale bar size = 10μm

YAP1 activation state influences its subcellular localization; inactivation causes redistribution of YAP1 from the nucleus to cytoplasm (Zhu et al., 2015b). Importantly, both extrinsic and intrinsic mechanisms can regulate YAP1 transcriptional activity (Hansen et al., 2015; Meng et al., 2016). To test whether cell fusion alters YAP1 activity, we looked at endogenous YAP1 localization in our VSV-G cell fusion system. Remarkably, we observed a shift from YAP1 being primarily in the nucleus in non-fused cells to being in the cytoplasm in fused cells (Figure 3C, top panel). Analyses of YAP1 localization at different time points revealed this occurred within 1 hr and was maintained at least up to 4 days after cell fusion (Supplementary Figure 3, Figure 3C-D, top panel).

Given this significant response, we investigated whether YAP1 was also primarily localized in the cytoplasm in physiological syncytia. For this, human trophoblasts purified from termed placenta were cultured and allowed to fuse during 2, 3 and 4 days (Kliman et al., 1986; Tang et al., 2011). Analyses of YAP1 localization revealed fused trophoblast contain predominately cytoplasmic YAP1 localization as we observed for our cell fusion model system (Figure 3C-D, bottom panel). Analysis of YAP1 localization in tissue sections from mouse skeletal muscle in developing mouse embryos (E10.5) revealed YAP1 localization was also primarily cytoplasmic within MF20 positive syncytia while it was mostly localized in the nucleus of PAX7 positive progenitors cells (Figure 3E, F) (Chal and Pourquie, 2017). This suggests YAP1 inactivation and cytoplasmic redistribution is a hallmark of fused cell systems that could be driving their differentiation state.

### Cell fusion promotes increased endocytosis altering plasma membrane surface area

We next evaluated whether specific changes in fundamental cellular features occurring during cell fusion were responsible for YAP1 inactivation and transcriptional reprogramming. Changes in shape are known to be governed by alterations in both volume and surface area (SA) (Harris and Theriot, 2018; Okie, 2013). Importantly, recent work has shown that YAP1 activity and cellular localization is altered through variations in volume and/or SA (Hong et al., 2017; Low et al., 2014; Nardone et al., 2017; Perez Gonzalez et al., 2018; Cai et al., 2019). Given our observation that fusing cells undergo a significant change in cell shape, we wondered whether this could be related to changes in YAP1 localization upon cell fusion.

To measure SA during cell fusion, cells expressing a fluorescence PM marker were used to create surfaces using the 3D rendering software Imaris. Surprisingly, immediately after cells start to fuse membrane protrusions are observed and the SA starts to decrease (Figure 4A, Supplementary Video 6). To measure the extent by which cell fusion alters SA and volume, we compared the SA and volume of cells before (Pre-Fusion) and 30 minutes after cell fusion (Post-Fusion) (Figure 4B). We observed a 15% decrease in PM SA after cell fusion was induced, while volume only modestly changed (Figure 4B-D). This suggests that upon fusion the newly formed syncytium activates adaptive responses that modulate its plasma membrane SA while keeping its volume relatively constant.

**Figure 4.**
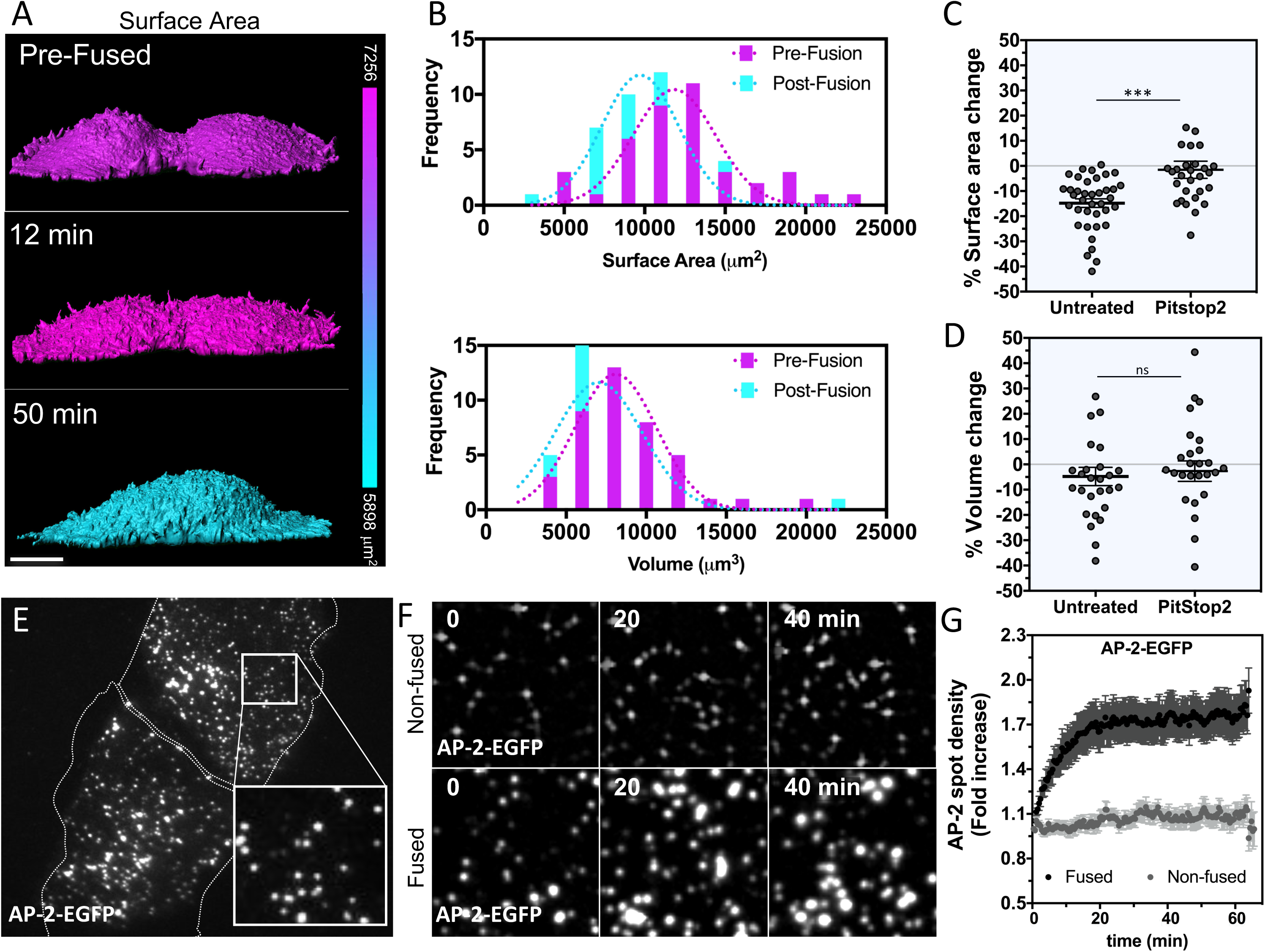
Remodeling upon cell fusion reduces the plasma membrane surface area by increasing endocytosis. (A) SUM-159 cells expressing VSV-G and a PM marker (CAAX-EGFP) were imaged by confocal microscopy as cells fused (Z-stacks were used to generate three-dimensional models of cells). The surface of fusing cells is colored based on the surface area as measured by the IMARIS surface tool (See also Supplementary Video 6). (B) The frequency distribution of surface area and volume of pairs of cells within 30 minutes of cell fusion. (C and D) The percentage change in plasma membrane surface area (C) and cell volume (D) 30 minutes after cell fusion with or without PitStop2 treatment to inhibit endocytosis. (E and F) CRISPR-Cas9 gene edited SUM-159 cells expressing endogenous AP-2 –EGFP and transfected with VSV-G, were induce to fused, and the density of AP-2 –EGFP at the plasma membrane was measured overtime by TIRF microscopy (using 100μm^2^ cropped images, such as the inset in E). Representative images of AP-2 –EGFP puncta at the indicated time points in non-fusing (F, upper) and fusing cells (F, lower). (G) The fold increase in AP-2 –EGFP puncta density was measured every minute for 1 hr (See also Supplementary Video 7). Means ± SEM are shown, (***) P = 0.0004, (no significant = ns) P > 0.05, Scale bar size = 10μm

The role of endo-exocytic pathways in the regulation of PM SA has been well documented (Morris and Homann, 2001). Therefore, our finding showing a decrease in PM SA suggests that upregulation of endocytosis might be mediating PM internalization. To test this hypothesis, the plasma membranes of fusing cells were labeled with a lipid fluorescence dye (DiD) and the internalization of vesicles after cell fusion was monitored. We observed that upon cell fusion the number of internalized vesicles increases (Supplementary Figure 4A). This supports the idea that upregulation of endocytosis during cell fusion leads to a reduction in PM SA, which can potentially influence YAP1 cellular distribution and transcriptional reprogramming.

### Up-regulation of CME is necessary for surface area regulation and YAP1 inactivity during cell fusion

Prior work has shown that clathrin-mediated endocytosis (CME) can regulate the SA of cells that are undergoing cell division (Aguet et al., 2016; Boucrot and Kirchhausen, 2007; Tacheva-Grigorova et al., 2013). To determine whether CME is also involved in the regulation of PM SA upon cell fusion, we assessed the frequency of clathrin coated pits by TIRF microscopy. Analyses of the steady-state levels of PM clathrin-heavy-chain (CHC) and the alpha-subunit of the clathrin adaptor AP-2 show CME increases after cell fusion (Supplementary Figure 4B). To accurately measure the degree by which CME augmented after the formation of fusion pores (T= 0 minutes, Supplementary Video 7), we used a gene edited cell line expressing endogenous levels of fluorescently labeled AP-2 (Kural et al., 2015) (Figure 4E). Consistent with upregulation in CME, there was a 75% increase in AP-2 spot density 20 minutes after cell fusion, whereas non-fused cells didn’t show any change in the levels of AP-2 spot density (Figure 4F, G). Furthermore, pre-treatment of fusing cells with the inhibitor of endocytosis, PitStop2, blocked the reduction in SA, suggesting that active CME is required for proper control of the PM SA during cell fusion (Figure 4C).

Given the significant effect of endocytosis in reducing the PM SA early after cell fusion, we asked whether this was the upstream regulatory mechanism leading to YAP1 redistribution. To address this, we examined whether inhibiting CME blocks YAP1 cytoplasmic retention. Remarkably, pre-incubation of fusing cells with 2 different inhibitors of endocytosis (PitStop2 and Dynasore) strongly decreased YAP1 redistribution into the cytoplasm (Figure 5A, B). Furthermore, expression of the CME specific inhibitor AP180-C showed a higher impact on preventing YAP1 cytoplasmic redistribution, demonstrating the specific role of CME in YAP1 regulation during cell fusion (Figure 5A, B) (Zhao et al., 2001).

**Figure 5.**
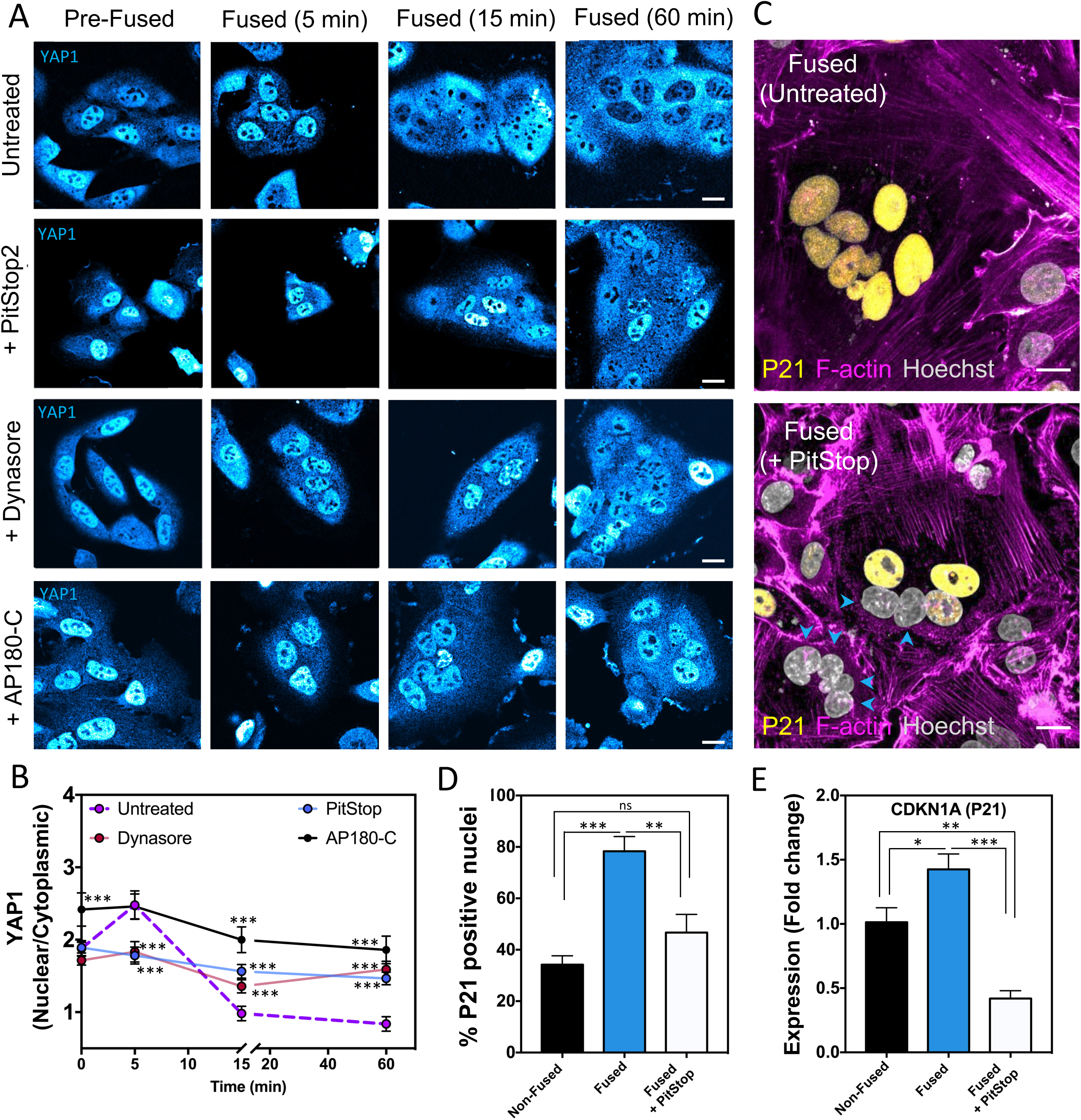
YAP1 inhibition and cell-cycle arrest after cell fusion depend on active clathrin-mediated endocytosis. (A) SUM-159 cells, transfected with VSV-G, were incubated with inhibitors of endocytosis (PitStop2 or Dynasore) or transfected with a dominant negative form of the clathrin adaptor AP-180 (AP180-C), a specific inhibitor of CME. Cells were then induced to fuse, fixed at indicated time points, immunostained with anti-YAP1 antibody and imaged by confocal microscopy to determine the YAP1 localization. (B) The ratio of nuclear to cytoplasmic YAP1 was measured at each timepoint after fusion in control (untreated) or endocytosis inhibited cells (P values represent the differences between the control and each endocytic inhibitor). (C) Immunofluorescence staining of fused cells to identify P21 positive nuclei in untreated and Pitstop2 treated cells. (D) The percentage of P21 positive nuclei measured in immunostained cells (cells were fixed 24hr after fusion) and (E) the expression of the P21 transcript (CDKN1A) measured by qRT-PCR is compared in non-fused cells, fused cells, and fused cells treated with PitStop2. Arrow heads points at nuclei within fused cells negative for P21. Means ± SEM are shown, (***) P < 0.0001, (**) P < 0.001, (*) P < 0.05 (no significant = ns) P > 0.05, Scale bar size = 10μm

Since inhibition of CME strongly blocks YAP1 nuclear exclusion, we reasoned that upregulated endocytosis might be a required step for transcriptional reprogramming after cell fusion. Importantly, prior work has shown that down-regulation of YAP1 can lead to cell-cycle arrest and the up-regulation of P21 (Liu et al., 2017). To test whether inhibition of endocytosis, which leads to YAP1 nuclear retention, blocks cell-cycle arrest we measured the levels of P21 in fused cells pre-treated with PitStop2. We found that fused cells treated with PitStop2 have fewer P21 positive nuclei than untreated fused cells (Figure 5C, D). Consistent with these findings, qRT-PCR analyses revealed that inhibition of endocytosis also reduces the expression levels of P21 (CDKN1A transcript) in fused cells (Figure 5E). This demonstrates that active endocytosis acts as a cell-intrinsic cue leading to YAP1 nuclear exclusion and the expression of cell-cycle arrest genes.

### Increased CME upon cell fusion results in transient glucose transporter depletion and acute energy stress

Next, we addressed the CME downstream intracellular mechanism leading to YAP1 cytoplasmic redistribution and transcriptional reprogramming. Active endocytosis not only regulates PM SA in cells, but is also essential for controlling the surface distribution of a wide-range of membrane proteins including surface receptors and transporters (Antonescu et al., 2014; Bokel and Brand, 2014; Weinberg and Puthenveedu, 2018). Recent studies have suggested that surface expression of glucose transporters and the levels of cytoplasmic glucose can control YAP1 localization and activity (Santinon et al., 2015; Zhang et al., 2018). Therefore, one possibility is that increased CME upon cell fusion changes the distribution of glucose transporters leading to YAP1 nuclear exclusion. To test this possibility, we examined whether cell fusion triggers the internalization of glucose carriers.

The most widely expressed glucose transporter isoform, Glut1 (Augustin, 2010; Mueckler and Thorens, 2013) localizes largely to the plasma membrane in control cells (non-fused). In contrast, we observe an accumulation of Glut1 positive internal structures approximately 5 minutes after cell fusion was triggered. These internal structures start decreasing after 15 minutes and almost completely disappear after 60 minutes, suggesting Glut1 transporters are actively recycled back to the PM (Figure 6A, top panel and insets). Glut1 can be internalized by both CME and clathrin-independent endocytic (CIE) pathways (Antonescu et al., 2014). To test whether inhibition of CME affects Glut1 internalization upon cell fusion, we analyzed the localization of Glut1 after fusion of cells previously transfected with AP180-C. Consistent with a specific role for CME during cell fusion, expression of AP180-C blocks the internalization of Glut1 in fused cells (Figure 6B, lower panel and insets). Furthermore, an alternative localization analysis of the clathrin-dependent cargo transferrin receptor (TfR-GFP), revealed TfR-GFP is also internalized 5 minutes after fusion (3 fold) and, similar to glucose transporters, is partially recycled back to the PM after 60 minutes (Supplementary Figure 5). As expected, treatment with PitStop2 impaired TfR-GFP internalization (Supplementary Figure 5). Conversely, CD147 and CD98, two amino acid transporters known to be regulated by CIE, did not significantly internalize after cell fusion (Supplementary Figure 6). These results demonstrate that upregulation of CME during cell fusion can directly and specifically influence the surface levels of glucose transporters thus acutely modulating the PM landscape of the new syncytium.

**Figure 6.**
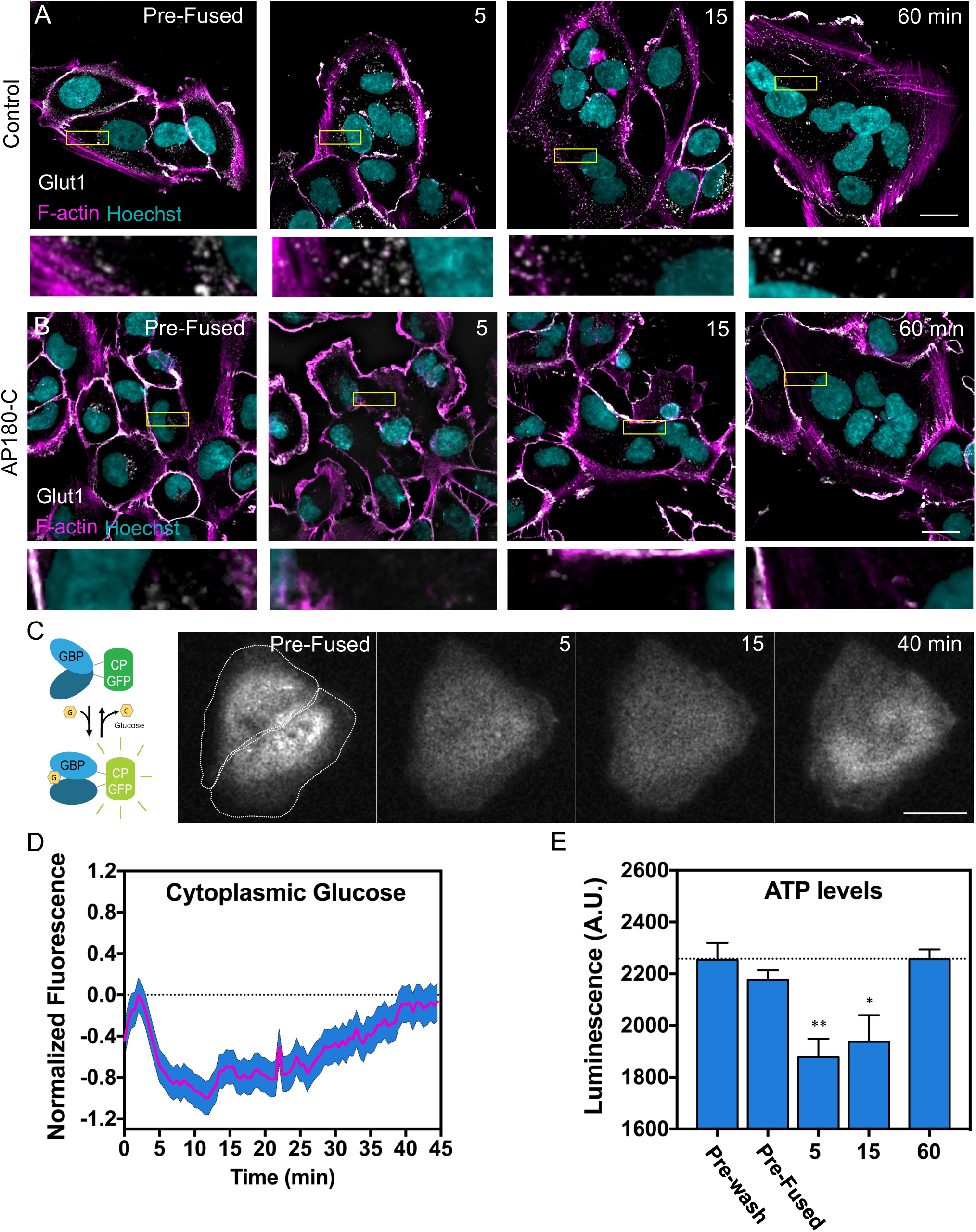
Cell fusion leads to a low energy state through transient, CME-dependent glucose channel internalization. (A and B) SUM-159 cells expressing VSV-G (A) or VSV-G and AP180-C (B) were induced to fused and then fixed at indicated time points. Anti-Glut1 antibodies were used to assess the subcellular localization of Glut1. Insets depict a region in the cytoplasm at each time point. (C) The glucose biosensor, (depicted in left panel, GBP = Glucose Binding Protein) fluoresces when glucose is bound. The intensity of the cytoplasmic biosensor fluorescence was monitored as cells stably expressing VSV-G were induced to fuse. (D) Relative cytoplasmic glucose levels were measured over-time in fusing cells. To calculate the relative changes the fluorescence intensity in fusing cells was normalized to the intensity of control non-fusing cells. (E) VSV-G transfected SUM-159 cells were fixed before fusion, or at the indicated time after fusion was induced. Relative cytoplasmic ATP levels were measured using a luciferase based assay. Means ± SEM are shown, (**) P < 0.003, (*) P < 0.03, Scale bar size = 10μm

The down-regulation of glucose transporters leads to low levels of cytoplasmic glucose and could lead to energy stress (Klip et al., 1994; Sasson et al., 1997). To measure the levels of glucose after cell fusion, we utilized a glucose biosensor that increases or decreases its fluorescence intensity depending on whether it is in its bound or unbound state, respectively (Figure 6C) (Keller and Looger, 2016). We measured a maximum decrease in cytoplasmic glucose approximately 12 minutes after induction of cell fusion followed by the recovery of cytoplasmic glucose after 40 minutes (Figure 6D). This result is consistent with our finding showing a fast internalization of glucose transporters within 5 minutes after cell fusion followed by recycling (Figure 6A). Parallel to the results with the glucose biosensor, luciferase-based ATP measurements detected a decrease in ATP levels 5 minutes after cell fusion followed by its recovery after 60 minutes (Figure 6E). This demonstrates that cell fusion induces an acute low energy state caused by fast internalization of glucose channels dependent on CME.

### AMPK acts as a downstream effector of cell fusion-induced structural changes to initiate gene reprogramming

When cells are depleted of ATP, the AMP-activated protein kinase (AMPK) is phosphorylated and activated (Hardie, 2018). AMPK is the master regulator of glucose metabolism and has the ability to sense the cytoplasmic AMP/ATP ratio. In addition, AMPK activity is important for cell differentiation-promoting processes including the regulation of YAP1 (Santinon et al., 2016). To test whether the low energy state induced upon cell fusion leads to activation of AMPK, we determined the levels of phosphorylated AMPK (P-AMPK) by biochemical analyses. We found a 2-fold increase in P-AMPK levels 5 minutes after cell fusion that is maintained for up to 60 minutes decreasing thereafter (Figure 7A-B). This is consistent with our measurements of GLUT1 internalization and recycling (Figure 6A). Since the low energy state induced by cell fusion is mostly caused by the transiently increased internalization of glucose channels and reduction of cytoplasmic glucose, we tested whether inhibition of endocytosis blocks the activation of AMPK. For this we pre-treated fusing cells with PitStop2 and measured the levels of P-AMPK. Similar to pre-treatment with the AMPK inhibitor, compound C, inhibition of endocytosis by PitStop2 blocked AMPK phosphorylation (Figure 7B). This demonstrates that activation of AMPK downstream of PM remodeling events is triggered by cell fusion.

**Figure 7.**
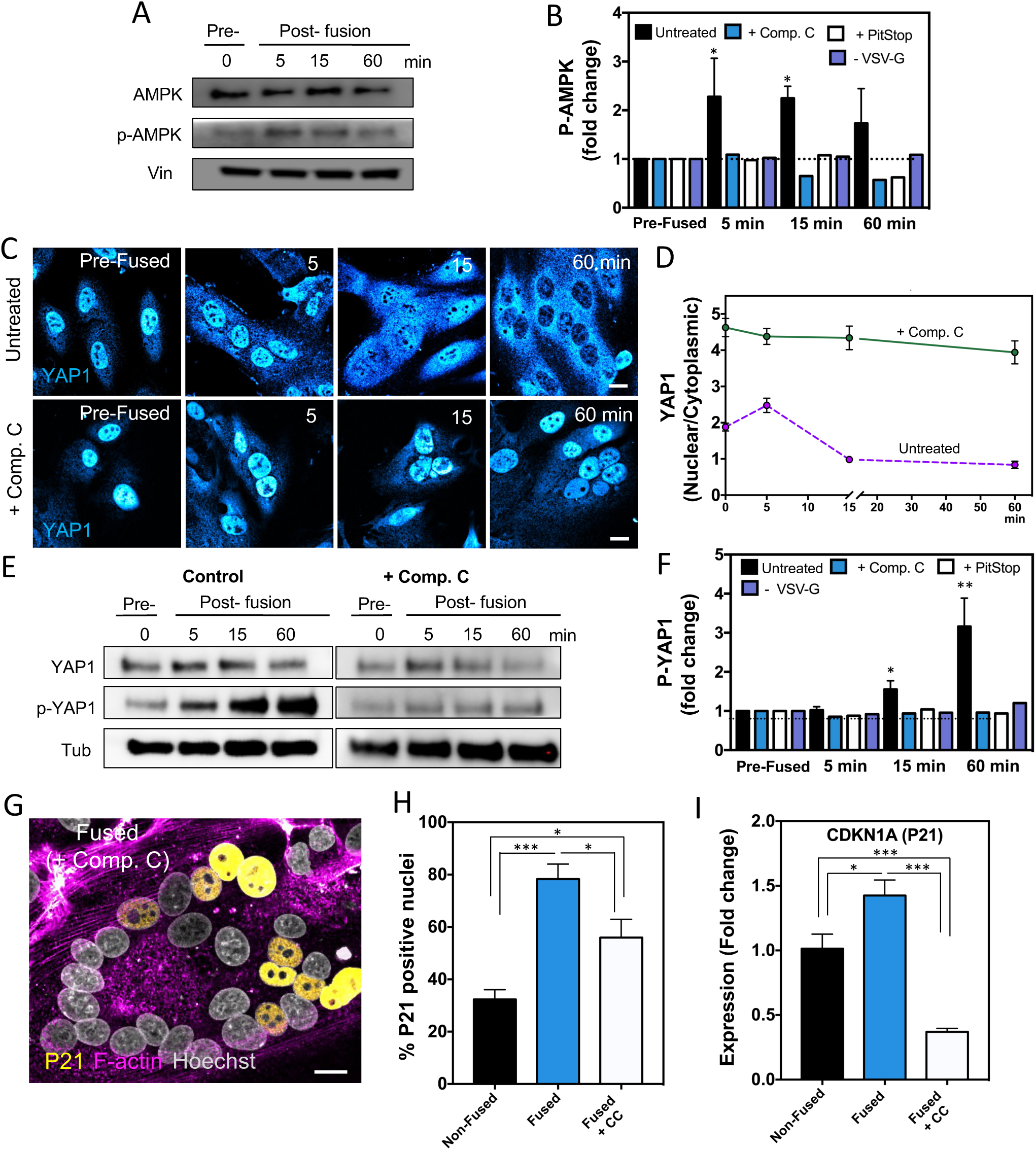
AMPK is activated upon cell fusion and AMPK inhibition impairs both YAP1 cytoplasmic localization and cell-cycle arrest. (A) Cell lysates were made from VSVG-transfected SUM-159 cells before (Pre-fusion; 0 min) or 5, 15, and 60 minutes after fusion. The levels of total and phosphorylated AMPK were determined by western blot. Vinculin (Vin) was used as a loading control. (B) The fold change of p-AMPK in lysates at the indicated times after fusion was calculated relative to the non-fused cells. Cells transfected with VSV-G were untreated, or treated with AMPK or endocytosis inhibitors (compound C or PitStop2). In addition, p-AMPK was quantified in untreated cells that lacked VSV-G (-VSV-G, unable to fuse). (C and D) SUM-159 cells incubated with or without the AMPK inhibitor compound C, were induced to fuse and then fixed at indicated time points and immunostained with an anti-YAP1 antibody (C). The YAP1 nuclear to cytoplasmic ratios in both treated and untreated samples were quantified (D). (E) Cell lysates from VSV-G transfected cells, either untreated or incubated with compound C, were prepared at the indicated times after fusion. Total YAP1 and YAP1 phosphorylation (S127) were assayed by western blot. (Tubulin (Tub) was used as a loading control). (F) The fold changes of p-YAP1 were calculated in untransfected (-VSV-G) cells, and in VSVG-transfected cells that were untreated or incubated with either compound C or PitSop2. (G and H) Nuclear P21 was visualized in compound C treated cells fixed and immunostained 24hr after fusion. (H) The percentage of P21 positive nuclei in compound C treated cells is compared to the non-fused and untreated cells. (I) CDKN1A (P21) expression levels in compound C treated and untreated cells were measured by qRT-PCR. Means ± SEM are shown, (***) P < 0.0001, (**) P < 0.001, (*) P < 0.05 (no significant = ns) P > 0.05, Scale bar size = 10μm

**Figure 8.**
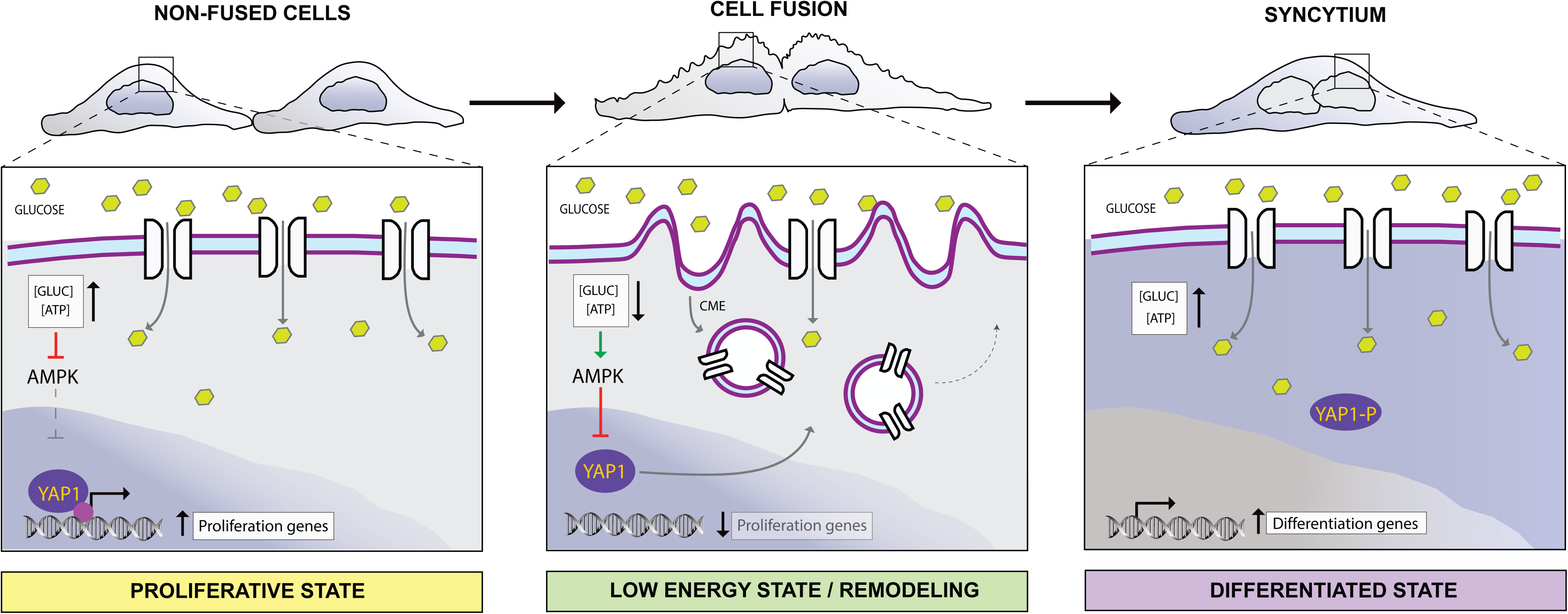
Structural remodeling upon cell fusion leads to endocytosis and AMPK-dependent YAP1 inhibition, which drives cell-cycle arrest and promotes a differentiated-like state. In proliferating cells, nuclear YAP1 promotes expression of genes to support the proliferative state. The acute structural remodeling upon cell fusion, including endocytosis of glucose transporters, causes transient changes in cell energetics (decreased cytoplasmic glucose and ATP) and AMPK activation that lead to persistent retention of YAP1 in the cytoplasm. As a result, fused cells exit the cell cycle and transcripts that promote cell differentiation are generated.

Prior work has shown that AMPK can negatively regulate YAP1 when cells experience a low energy environment (DeRan et al., 2014; Mo et al., 2015; Wang et al., 2015). We reasoned that AMPK activation could link the structural and energetic changes occurring during cell fusion to the downstream inhibition of YAP1 in fused cells. To test whether YAP1 nuclear exclusion upon cell fusion requires active AMPK, we treated fusing cells with compound C for 3hr and measured endogenous YAP1 distribution. Remarkably, inhibition of AMPK completely blocked YAP1 re-localization to the cytoplasm showing a predominant nuclei localization (Figure 7C, D). In addition, biochemical analysis of YAP1 phosphorylation confirmed that fusing cells treated with compound C or endocytic inhibitors have lower levels of inactive P-YAP1 compared with non-treated cells (Figure 7E, F). To test whether inhibition of AMPK (which also leads to YAP1 nuclear retention) blocks cell-cycle arrest, we measured levels of P21 in fused cells pre-treated with compound C. Similar to PitStop2 treatment, we found that fused cells treated with compound C have fewer P21 positive nuclei compared to untreated fused cells (Figure 7G, H). Furthermore, qRT-PCR analysis of compound C treated cells indicated that AMPK inhibition decreases the expression of P21 (CDKN1A transcript) (Figure 7I). Altogether, these results demonstrate cell fusion leads to an acute activation of AMPK driving YAP1 nuclear exclusion and supporting a differentiated state independent from tissue specific cues.

## Discussion

Cell fusion *in vivo* involves fast rearrangements of cell structural features, and long-term reprogramming of gene expression and differentiation. How the initial structural changes that accompany cell fusion are integrated to alter cell fate determination are still unknown. This gap in understanding stems from the challenges in dissecting the intrinsic contribution of cell fusion from extrinsic differentiation signals within tissues. Here, we have employed a viral protein (VSV-G) to induce fusion of culture cells in the absence of tissue specific differentiation cues. We identified several mechanisms regulating key transitions that cause fused cells to stop proliferating and diverts them toward a differentiation-like state. We observed that cell fusion shifted the surface area-to-volume ratio and triggered CME to remove excess membrane. As a result, glucose transporters, including Glut1, were temporarily internalized leading to reduced cytoplasmic glucose and ATP levels. The transient drop in cytoplasmic ATP activated AMPK, which promoted the phosphorylation of YAP1, shifting its localization from the nucleus to the cytoplasm. Consequently, the transcription of several genes involved in cell proliferation dropped, and genes involved in cell-cycle arrest and differentiation were expressed. Furthermore, we showed that disruption of either CME or AMPK activation prevents subsequent parts of the pathway. Importantly, the key feature – exclusion of YAP1 from the nucleus – is recapitulated in differentiated placenta cells and developing muscle tissue. These results demonstrate that physical and structural changes upon cell fusion are sufficient to trigger an intrinsic response that directs fused cells to transition into a new cellular state.

CME is the primary endocytic pathway regulating plasma membrane SA during fundamental processes such as cell division (Aguet et al., 2016; Tacheva-Grigorova et al., 2013). How does cell fusion induce increased CME? Prior work has demonstrated that the level of tension of the plasma membrane regulates the equilibrium between exocytosis and endocytosis (Diz-Munoz et al., 2013; Morris and Homann, 2001; Pontes et al., 2017; Boulant et al., 2011; Ferguson et al., 2017; Heuser and Anderson, 1989). Specifically, exocytosis is stimulated by high membrane tension to add more membrane, while endocytosis responds to low membrane tension in order to remove it (Diz-Munoz et al., 2013). We speculate, therefore, that excess plasma membrane at the interface where two cells have fused is sensed as low membrane tension, and that this helps trigger the increased CME during cell fusion.

AMPK has been shown to influence endocytosis and exocytosis of membrane proteins including ion channels and nutrient transporters (Foley et al., 2011; Gusarova et al., 2009; Puljak et al., 2008; Samovski et al., 2012; Walker et al., 2005; Weisova et al., 2009; Wu et al., 2013; Xi et al., 2001). Under energy stress, AMPK can increase the surface expression of Glut1 and Glut3 by promoting exocytosis (Cura and Carruthers, 2012; Weisova et al., 2009). We show Glut1 is rapidly internalized 5 minutes after the onset of cell fusion and eventually recycled back to the plasma membrane. It is possible that activated AMPK promotes the recycling of glucose transporters to restore glucose levels. This is consistent with our measurements of cytoplasmic glucose using a glucose biosensor showing that both the decrease and recovery of cytoplasmic glucose coincides with the internalization and recycling of glucose transporters.

YAP1 can directly control cell renewal by localizing to the nucleus where it promotes proliferative transcriptional programs (Hansen et al., 2015; Pan, 2010; Zhu et al., 2015a). In non-fusing cells, cytoplasmic localization of YAP1 has been strongly associated with differentiation of multiple cell types including keratinocytes, adipocytes and neurons (Panciera et al., 2017). However, in fusing cells, little is known about YAP1 distribution and its role during differentiation. Prior work *in vitro* suggested that myoblast differentiation leads to YAP1 cytoplasmic localization and reduced expression (Fischer et al., 2016; Watt et al., 2010). Here we demonstrated that the nuclear to cytoplasmic ratio of YAP1 *in vivo* decreased during murine muscle fusion and differentiation (Figure 3E). In addition, we demonstrated that YAP1 localization in human syncytiotrophoblasts and non-fused human trophoblasts paralleled the YAP1 nuclear to cytoplasmic shift observed in the VSV-G mediated fusion system (Figure 3C-D). These results establish that YAP1 cytoplasmic redistribution is an essential step and potentially a hallmark of syncytia differentiation triggered by cell fusion.

In summary, our data provides new insight into how intrinsic cellular changes arising from the fusion of two or more cells induces a differentiation-like state. We describe a structural-to-transcriptional signaling pathway mediated by an endocytosis-AMPK-YAP1 axis that links membrane remodeling and cellular bioenergetics to transcriptional reprogramming in response to cell fusion. In this pathway, AMPK plays a central role in sensing the transient reduction in cytoplasmic glucose and ATP caused by modifications in the plasma membrane landscape through increased CME. It then converts these changes into an adaptive response that inhibits YAP1 activity, inducing cell-cycle arrest and supporting the expression of differentiation-related genes. Hence, disabling either CME or AMPK by genetic or pharmacologic approaches hinders a transcriptional program towards differentiation during cell fusion. The broad transcriptional changes revealed by RNA-Seq, which include many transcripts not regulated by YAP1, suggest this pathway is likely integrated with other cellular signaling routes to achieve the cell state transition. Future work will be needed to reveal how this structural-to-transcriptional signaling pathway synergizes with additional environmental cues to accomplish functionally differentiated syncytia within tissues.

## Acknowledgments

We thank Andrew Lemire, Jingqun Ma, and Kshama Aswath for help with RNA-seq data generation; Kevin McGowan for help with the qRT-PCR data generation; Tomas Kirchhausen for provide with the SUM-159 and SUM-159-AP2-EGFP cells; Jacob Keller and Loren Looger for sharing the iGlucoSnFR.mRuby2 plasmid; Ulrike Heberlein and Erik Snapp for helpful discussion and comments on the manuscript. The work was supported by the Howard Hughes Medical Institute.

## Author Contributions

DF and JLS designed experiments. DF performed and analyzed *in vitro* experiments. DF and IEM performed and analyzed *in vivo* immunofluorescence experiments. DF and IEM analyzed RNAseq data. DF, KMM, ZT, HJK, and SMG performed human throphoblast purification and fusion experiments. DF and AW performed LLS microscopy experiments. DF, CMO, and JLS wrote the manuscript.

## Declaration of Interests

The authors declare no competing interests

## Supplementary Figure Legends

Supplementary Figure 1. **Intermixing of small cytoplasmic proteins and organelles during cell fusion.** (A) Diagram of VSV-G mediated cell fusion. Cells expressing VSV-G and different subcellular markers are rapidly washed (5-10 seconds) with Fusion Buffer to induce cell fusion. Fusion of cells occurs 30-60 seconds after wash. (B) Exchange of a small fluorescent cytoplasmic protein (mEmerald) between Donor and Receiver cells during cell fusion (observed as a measured decrease in fluorescence intensity in Donors and an increase in fluorescence intensity in Receivers). Note that, depending on the time of fusion, the rate of cytoplasmic mixing varies. Each curve is the average fluorescence of the cytoplasmic marker at 3 different ROIs within each cell. (C) Cells expressing either VSV-G or VSV-G and an ER marker (oxVenus-KDEL) were culture together and then fused. ER mixing was monitored as the ER marker spread into Receiver cells. (D) Cells expressing only VSV-G or VSV-G, a cytoplasmic marker (TagBFP2), and a mitochondria marker (Mito-EGFP) were culture together and then fused. Initial fusion is observed when the cytoplasmic marker enters the Receiver cells, and mitochondria mixing is determined by the gain of the labeled mitochondria. (E and F) To assess nuclear clustering, cells transiently co-transfected with VSV-G and the nuclear marker H2B-mCherry were imaged live during and after cell fusion and nuclei displacement and clustering were tracked (E). The movement of multiple nuclei simultaneously within the same syncytium is displayed relative to their individual distances from the nuclei cluster at different time points after cell fusion. Different tracks (nuclei) are depicted in different colors (See also Supplementary Video 5). Scale bars = 10 µm.

Supplementary Figure 2. **Cell division stops after VSV-G mediated cell fusion.** (A) Representative images of fused SUM-159 cells monitored for 36 hr after VSV-G mediated fusion. (B) 95% of fused cells no longer divided after cell fusion (N=40). Scale bars = 10 µm.

Supplementary Figure 3. **YAP1 cytoplasmic localization persists for hours after cell fusion.** (A) Non-fused or Fused SUM-159 cells were fixed and immunostained using antibodies against YAP1 at different times during and after fusion, then imaged by confocal microscopy. (B) The YAP1 nuclear to cytoplasmic ratios were measured (N=32-40). Scale bars = 10 µm.

Supplementary Figure 4. **Clathrin-mediated endocytosis increases after cell fusion.** (A) VSV-G transfected SUM-159 cells were stained with a lipid fluorescent dye (DiD) to monitor internalizing vesicles upon cell fusion (PM in magenta, mask detecting internalized vesicles in green). (B) Immunofluorescence TIRF-microscopy of SUM-159 cells double labeled with antibodies to the clathrin heavy chain (CHC) and the alpha-subunit of the clathrin adaptor protein 2 (AP-2). The number of positive endocytic sites (spots) per 100 um^2^ were quantified before (Non-Fused) and after fusion (15, 30 min).

Supplementary Figure 5. **The clathrin-dependent cargo, transferrin receptor, is internalized upon cell fusion.** Cells transfected with both VSV-G and the transferrin receptor (TfR-EGFP) were culture with or without PitStop2, and then induced to fused. Internalization of TfR-EGFP was quantified using a bromophenol blue (BPB) quench-assisted localization assay. The fluorescence intensity before and after surface quenching with BPB were quantified at different time points before or after fusion (0, 5,15, 60 min). The ratio of Total to Internal fluorescence is displayed (see Methods).

Supplementary Figure 6. **Clathrin-independent endocytic cargo does not internalize upon cell fusion.** (A) SUM-159 cells expressing VSV-G were induced to fused and then were fixed at different time points. Anti-CD98 (A) and Anti-CD147 (B) antibodies were used to assess the subcellular localization of these clathrin-independent endocytic cargos during cell fusion. No detectable internalization of these cargos was observed. Scale bars = 10 µm.

## Supplementary Video Legends

Supplementary Video 1. Cytoplasmic mixing after cell fusion of SUM-159 cells. Related to Figure 1A and B.

Supplementary Video 2. Lattice light sheet microscopy of fusion pore formation and plasma membrane remodeling after cell fusion. Related to Figure 1C.

Supplementary Video 3. Lattice light sheet microscopy of fusion pore formation and plasma membrane remodeling after cell fusion (orthogonal view). Related to Figure 1C (lower panel) and D.

Supplementary Video 4. Remodeling of the actin cytoskeleton during cell fusion. Cells expressing both VSV-G and lifeact-EGFP (F-actin) were imaged and the dynamics of filamentous actin were monitor after washing with Fusion Buffer. Images were acquired using AiryScan microscopy (Zeiss) at an acquisition rate of 1 frame every minute for 1hr. Color coding represent the Z-position of actin filaments. Scale bars = 10µm.

Supplementary Video 5. Cell fusion promotes nuclei clustering. Related to Supplementary Figure 1.

Supplementary Video 6. Cell fusion remodeling of the plasma membrane reduces total cellular surface area. Related to Figure 4.

Supplementary Video 7. The number of AP-2 positive endocytic sites increases in fusing cells but not in non-fusing cells. To determine T=0 in fusing cells, cytoplasmic mixing was monitored (TagBFP2, blue). Related to Figure 4.

## Methods

### Cell lines

The human mesenchymal triple-negative breast cancer-stem cell lines, SUM-159 and SUM-159-AP2-EGFP, were obtained as a gift from Tomas Kirchhausen. The U2OS and HEK 293T cell lines were obtained directly from ATCC (HTB96 and CRL11268, respectively). Cells were grown in a 37°C, 5% CO2 tissue culture incubator on tissue culture treated dishes in DMEM + 10% FBS, L-glutamine and antibiotics. Cell lines were passaged with 0.25% Trypsin EDTA.

### Plasmids

The pMD2.G VSV-G (#12259), H2B-mCherry (#20972), TfR-EGFP(#54278), and mEmerald (#53976) plasmids were obtained from Addgene. The plasmids for CAAX-EGFP(Farn-119), mTagBFP2-C1 (CV-261), Mito-EGFP (Clon-109) were obtained from the Michael Davison collection. iGlucoSnFR.mRuby2 was a gift from Loren Looger at Janelia Research Campus.

### Transfection

All cell lines were transfected with VSV-G and the corresponding expression vectors using Lipofectamine 3000 (ThermoFisher, Cat. # L3000015) following the manufacturer’s protocol.

### VSV-G mediated cell fusion

For live-cell fluorescence microscopy experiments, cell fusion was performed as described in (Feliciano et al., 2018). Briefly, cells transfected with VSV-G and cultured at 37°C, 5% CO2 were rapidly washed (5-10 seconds) with an isotonic low pH buffer (pH 5.5-6.0, Fusion Buffer 37°C) to induce fusion of the plasma membranes of two or more adjacent cells. After fusion was induced cells were rapidly returned to regular medium and imaged at 37°C, 5% CO2. For immunofluorescence analysis, cell fusion was induced sequentially at different times, and then all samples were fixed simultaneously in order to obtain the different time points of the fusion process. For immunoblot analysis, cells were fused and rapidly incubated at 37°C, 5% CO2. After the corresponding incubation time (0, 5, 15, 60 min) cells where placed at 4° C to slowdown intracellular processes and scrape off to be used for cell lysates.

### Human trophoblasts

Human trophoblasts, isolated from term placentas, were culture in coverslip chamber slides and allowed to fused for 48, 72, and 96 hrs as previously described (Kliman et al., 1986, Tang et al., 2011). After the corresponding time point, pre-fused cytotrophoblast and fused syncytiotrophoblast were fixed with 4% paraformaldehyde (PFA, EMS) and used for subsequent immunofluorescence experiments.

### Histology of mouse embryos

Pregnant CD-1 female mice were obtained from Charles River and the copulatory plug was labeled as day 0.5 dpc. At 10.5 dpc the mice were sacrificed by cervical dislocation and embryos were fixed for 3 hours in 4% paraformaldehyde (PFA, EMS). Immunofluorescence on sections was performed as follows: embryos were embedded in a 15% sucrose and 7.5% gelatin solution, frozen at −80°C and sectioned (25µm) using a Leica Cryostat. Primary antibodies were applied overnight in a PBS-Triton-FCS solution (PBS, 0.1%Triton X-100, 20%FCS). The slides were washed 3x for 10 min in PBS-Triton X-100 (0.1% Triton X-100), and secondary staining was performed in PBS-Triton-FCS containing Hoechst 33342 for 2 hours at room temperature. Slides were mounted with Fluoromount (Sigma, Cat. # F4680) and imaged using a Zeiss LSM880 confocal microscope. The following primary antibodies were used: anti-Pax7, Mouse IgG1, (DSHB, ID:AB528428), anti-MF20, Mouse IgG2b (DSHB, ID:AB2147781), and Anti-Yap1 (Cell Signaling). The following secondary antibodies were used: anti-mouse IgG2b Cy3 (Jackson Immunoresearch Laboratories), anti-mouse IgG2b A647 (Jackson Immunoresearch Laboratories), and anti-rabbit IgG A488 (Jackson Immunoresearch Laboratories). All animal experiments were conducted according to the National Institutes of Health guidelines for animal research. Procedures and protocols on mice were approved by the Institutional Animal Care and Use Committee at Janelia Research Campus, Howard Hughes Medical Institute.

### Immunofluorescence

SUM-159 cells plated on coverslip chambers (ThermoFisher, Cat. # 155379) were fused at their corresponding time points, fixed with 4% paraformaldehyde (PFA, EMS), and then permeabilized and blocked with blocking solution (0.5% TritonX-100, 10% BSA in PBS) for 1 hr. Primary antibodies were diluted in blocking solution and incubated overnight at 4°C. After 3x washes (10 min) with PBS, cells were incubated with Alexa Fluor-conjugated secondary antibodies. The following primary antibodies were used: Anti-Yap1 (Cell Signaling), Anti-p21 (Cell Signaling, Cat. # 2947S), Anti-pH3 (anti-Phospho-Histone H3 (Ser10); Cell Signaling, Cat. #3377), Anti-clathrin heavy chain (Abcam, Cat. # ab21679), Anti-AP-2 (Abcam, Cat. # ab189995), Anti-Glut1 (Abcam, Cat. # ab40084), Anti-CD98 (BioLegend, Cat. # 315602) and Anti-CD147 (BioLegend, Cat. # 306202). For imaging and quantification, at least 20 fields of view per well were randomly chosen by Hoechst 33342 nuclear staining (ThermoFisher, Cat. # 62249) and imaged by Zeiss LSM880 confocal or NIKON TIRF microscope. At least 3 different samples were quantified per treatment type at each respective time point.

### RNA Sequencing

To Isolate the RNA, fused or non-fused SUM-159 cells were lysed with TRIzol reagent (ThermoFisher, Cat. # 10296010). A second CHCl3 extraction was performed to increase RNA purity. Concentration and purity was determined by Nanodrop (ThermoFisher). RNA-seq libraries were made from 5 ng RNA per sample, using Ovation RNA-seq v2 (NuGEN) to make cDNA and Ovation Rapid DR Multiplex System (NuGEN) to make libraries according to the manufacturer’s protocol. ERCC Mix 1 spike-in controls (ThermoFisher) were added at 1e^-5^ final dilution. Libraries were pooled for sequencing on a NextSeq 550 instrument (Illumina) using 75 bp reads in paired-end mode. Sequencing reads were trimmed to remove TruSeq adapters using Cutadapt (DOI: https://doi.org/10.14806/ej.17.1.200), then were aligned to the human genome (Hg38) using STAR (DOI: 10.1093/bioinformatics/bts635). Transcript BAMs were generated by STAR and gene expression estimates were made using RSEM (DOI: https://doi.org/10.1186/1471-2105-12-323). Differential expression analysis was performed using EBseq (DOI: <u>10.18129/B9.bioc.EBSeq</u>) with FDR = 0.05. Gene enrichment analyses were performed using DAVID Bioinformatics Resources (DOI: https://david.ncifcrf.gov). We used ToppCluster and Cytoscape (DOI: https://toppcluster.cchmc.org/, https://cytoscape.org) to construct the subcategory network. Heatmaps of gene expression were generated using Morpheus (DOI: https://software.broadinstitute.org/morpheus/).

### RNA isolation and quantitative real-time PCR

Relative gene expression was determined using Taqman RNA-to-Ct 1-step kit (ThermoFisher) with TaqMan gene expression assays for CDKN1A (ThermoFisher). The RNA from fused or non-fused SUM-159 cells, that were treated or untreated with the endocytic inhibitor PitStop2 (Abcam, Cat. # ab120687) or the AMPK inhibitor compound C (Sigma, Cat. # P5499-5MG), was isolated using TRIzol (ThermoFisher, Cat. # 10296010). qRT-PCR reactions were initiated with 100 ng of RNA for each sample following manufacturers protocol. Data was acquired with a Roche480 light cycler. Samples were run on triplicate plates and their Ct values averaged. Relative quantitation was performed using Delta-Delta Ct method. Analysis was performed using phosphoglycerol kinase (PGK) or glyceraldehyde 3-phosphate dehydrogenase (GAPDH) as a reference gene.

### Immunoblot analysis

SUM-159 cells that had been scraped off plates as described above were lysed with lysis-buffer (50mM Tris-Cl, pH 7.5, 150mM NaCl, 0.1% SDS, 1mM DTT and 1mM EDTA) containing protease and phosphatase inhibitors (Sigma) for 30 min at 4° C. After lysing the cells, samples were centrifuged at top speed in a microcentrifuge and the supernatants were recovered for subsequent steps. Lysates were separated by SDS-PAGE on gels and transferred to PVDF. Blots were incubated with anti-YAP1 (Cell Signaling), anti-phospho-YAP1 (Cell Signaling (S127)), anti-AMPK (Cell Signaling, Cat. # 2532s), anti-phospho-AMPK (Cell Signaling, Cat. # 2531s), anti-Vinculin (Sigma, Cat. # V9131), and anti-Tubulin (Sigma). Secondary antibodies conjugated to HRP were used (ThermoFisher).

### ATP measurement

To determine the relative levels of cytoplasmic ATP at different time points during the fusion process, total cellular ATP was assayed using a luciferase based ATP determination kit following the manufacturer’s protocol (ThermoFisher).

### Surface Area and Volume measurements

To determine the surface areas and volume, SUM-159 cells transfected with VSV-G and a plasma membrane marker (CAAX-EGFP) were culture at low confluency and only pairs of cells adjacent to each other were imaged before and 30 minutes after cell fusion. PitStop2 (Abcam, Cat. # ab120687) treated and untreated cells were imaged to acquire Z-stacks using a Zeiss LSM880 confocal microscope. Determination of surface areas and volumes was achieved using the surface tool in the Imaris software (Bitplane).

### Glucose biosensor

Images of cells expressing VSV-G and the iGlucoSnFR.mRuby2 glucose biosensor were acquired on a Zeiss LSM880 confocal microscope immediately after cell fusion was induced. While the green channel was used to monitor cytoplasmic glucose levels during the fusion process, the red channel served to confirm that cells were stable during the experiment in the x-y and focal planes.

### TIRF microscopy

Live-cell and immunofluorescence imaging of clathrin and AP-2 at endocytic sites was performed by TIRF microscopy using a NIKON Eclipse Ti Microscope System equipped with an environmental chamber (temperature controlled at 37°C and CO2 at 5%), Apo TIRF 100X objective (NA 1.49), high-speed EM charge-coupled device camera (iXon DU897 from Andor), and NIS-Elements Ar Microscope Imaging Software.

### Bromophenol Blue (BPB) quenching assisted microscopy

HEK 293T cells transfected with both VSV-G and the transferring receptor (TfR-EGFP) were pre-cultured with or without PitStop2 (Abcam, Cat. # ab120687). Total fluorescence and BPB-quenched images were taken at different time points upon fusion (0, 5, 15, 60 min) and the Total / Internal fluorescence ratios were calculated. For the quenching step, Bromophenol blue (BPB) (Sigma, Cat. # B8026) was dissolved in phenol red-free DMEM containing 25 mM HEPES buffer (ThermoFisher, Cat. # 15630080) and applied at a final concentration of 2 mM to ensure instant and effective quench of EGFP fluorescence.

### Airyscan microscopy

To visualize changes in F-actin during cell fusion, U2OS cells stably expressing Lifeact-EGFP were transfected with VSV-G, washed with Fusion Buffer. Imaging was performed using a Zeiss LSM880 with Airyscan microscope with a plan-apochromatic 63x oil objective (NA=1.4). Images were processed with Airyscan processing in ZEN software (Zeiss).

### Lattice light-sheet microscopy

To monitor plasma membrane dynamics upon cell fusion, cells transfected with VSV-G and a plasma membrane marker (GPI-EGFP) were washed with Fusion Buffer before imaging on a custom-built lattice light sheet microscope (LLSM). Imaging was performed using 488nm excitation, and a multiband pass emission filter (NF03-405/488/532/635E, Semrock). Data were acquired by serial scanning of the entire fusing cells through the light sheet with a 3D imaging rate of 30s per volume. All acquired data were deconvolved by using a Richardson-Lucy algorithm adapted to run on a graphics-processing unit, using an experimentally measured PSF.

