## Supplementary figures and images for "Transcriptional reprogramming in fused cells is triggered by plasma-membrane diminution"

### Supplementary Figures

Supplementary Figure 1.

Feliciano *et al.*

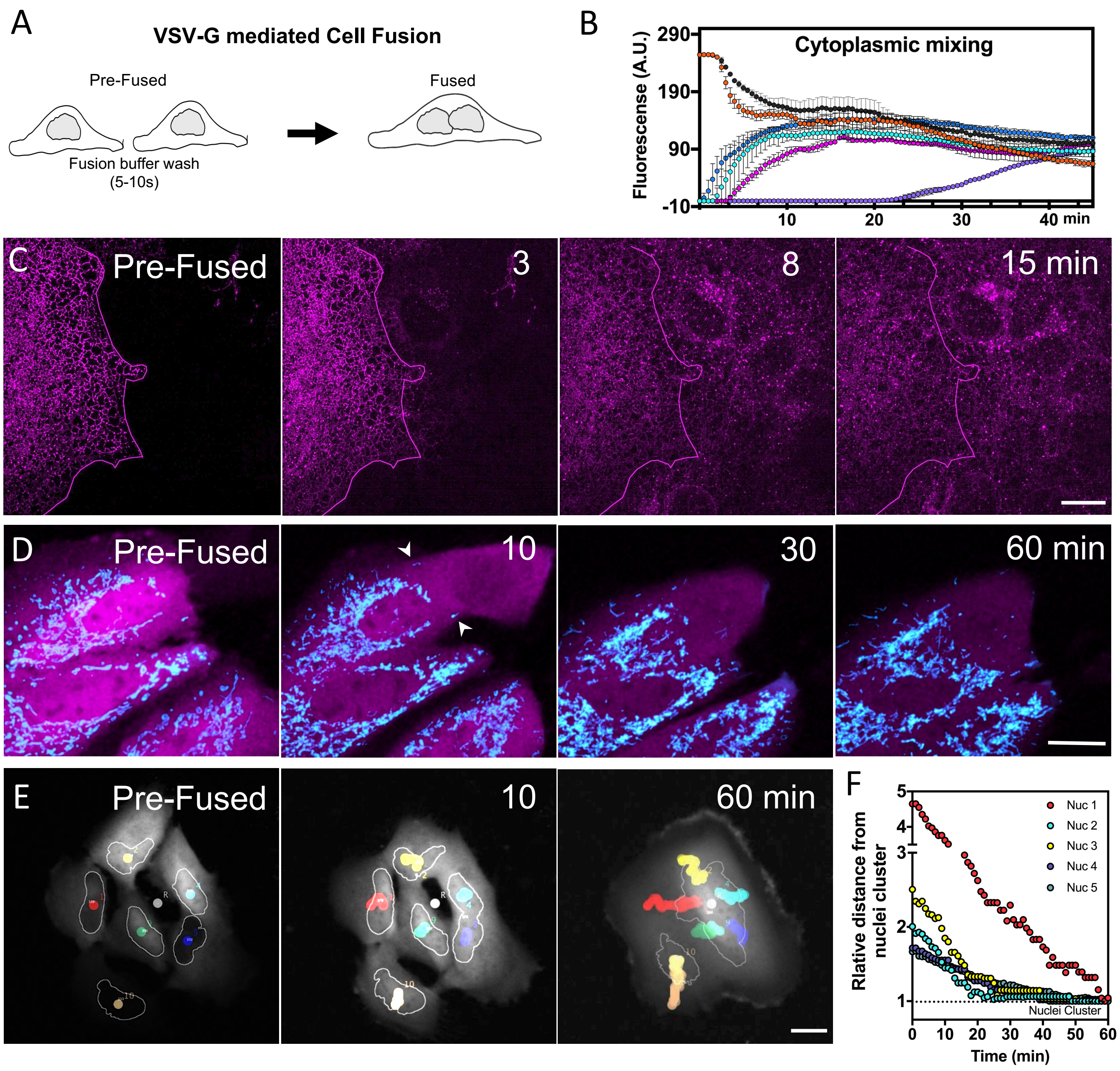

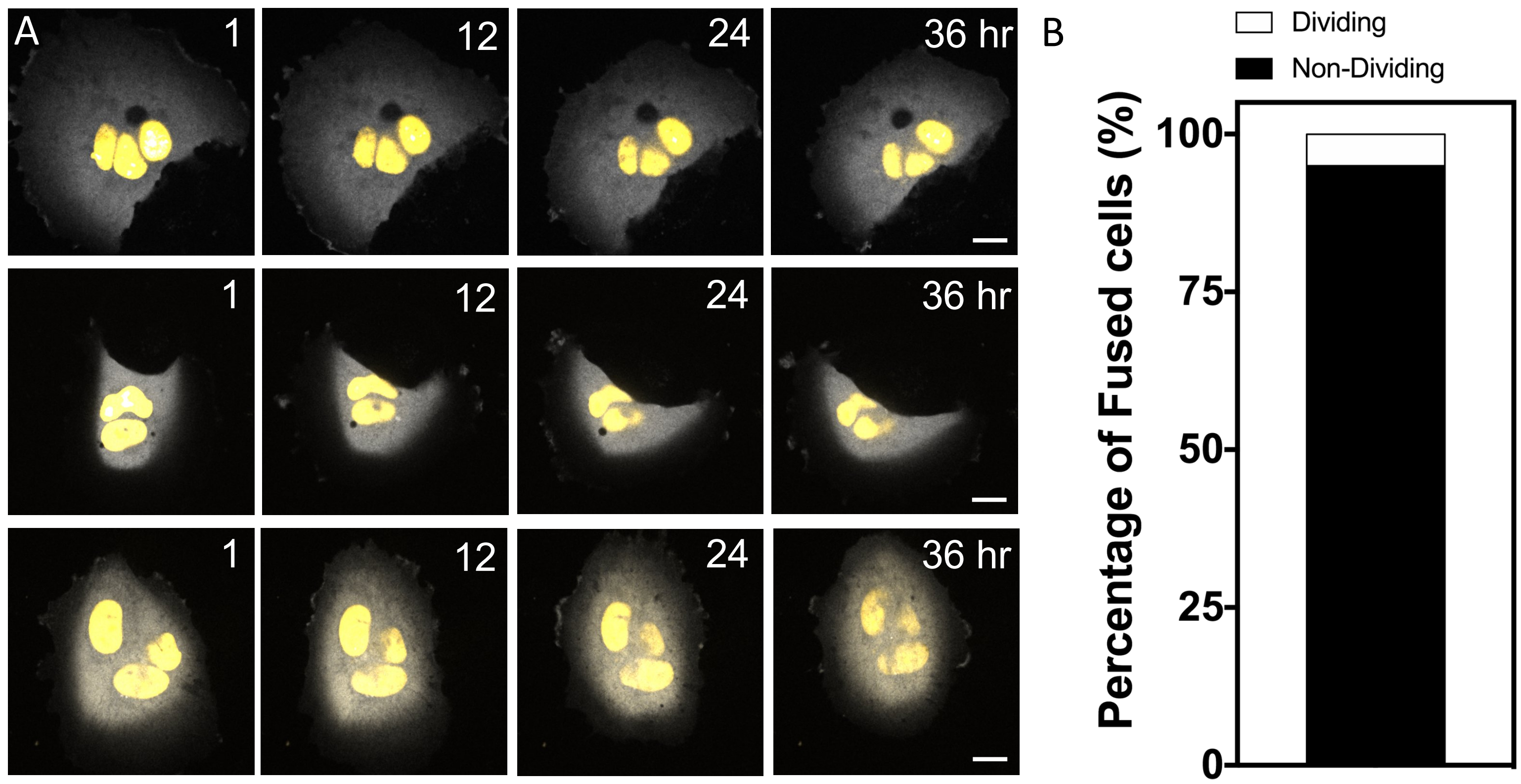

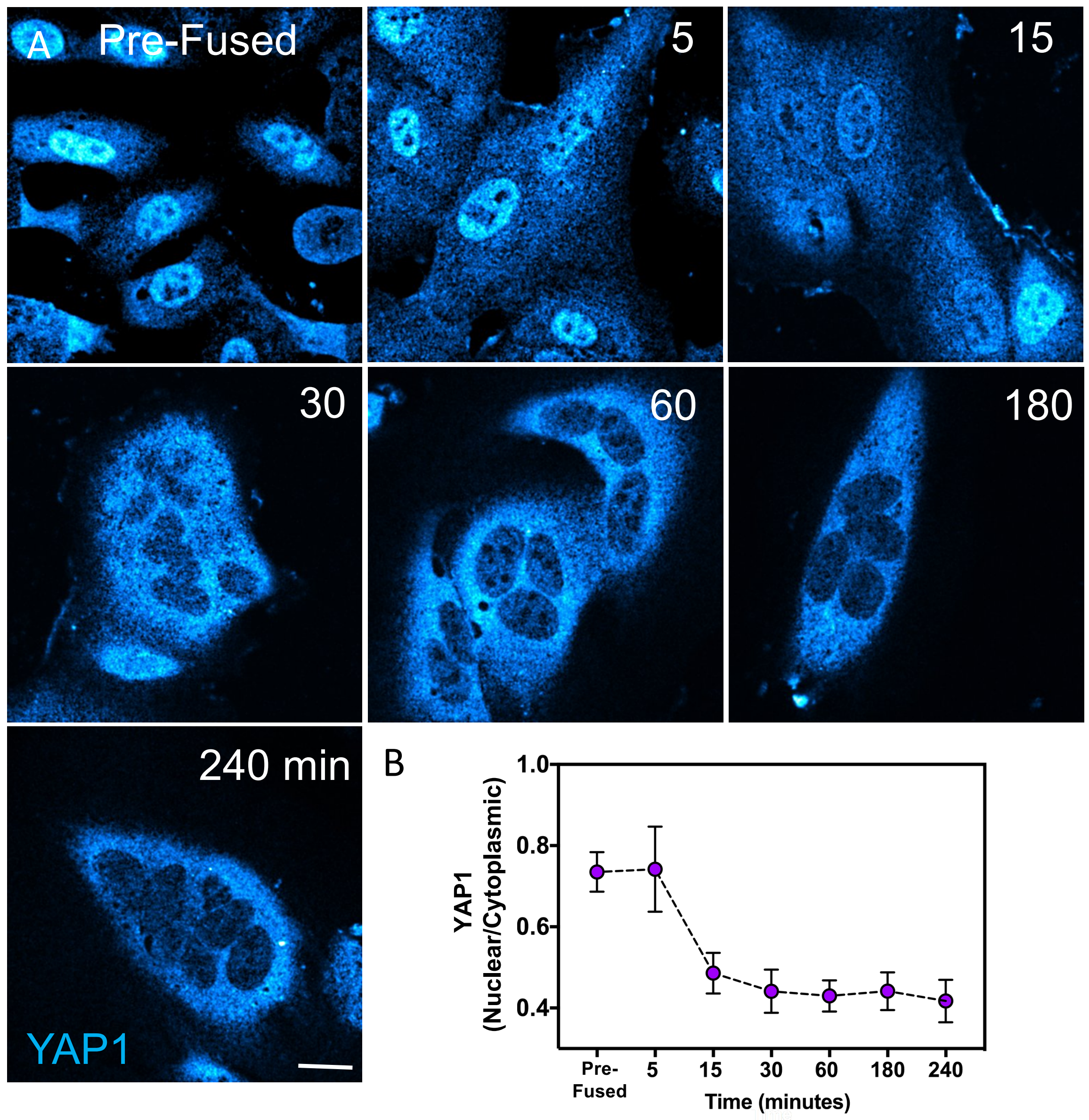

Supplementary Figure 4. Feliciano *et al.*

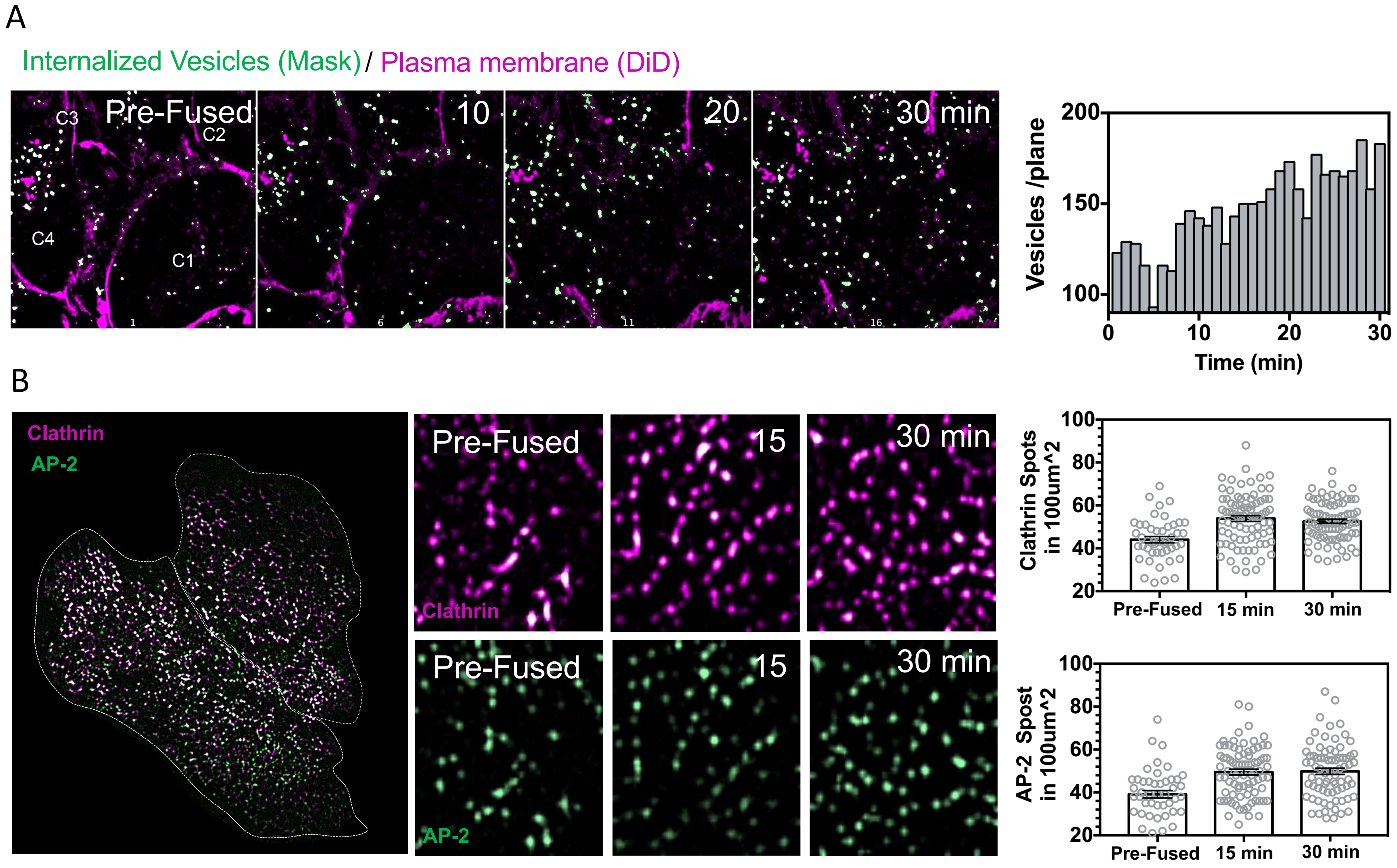

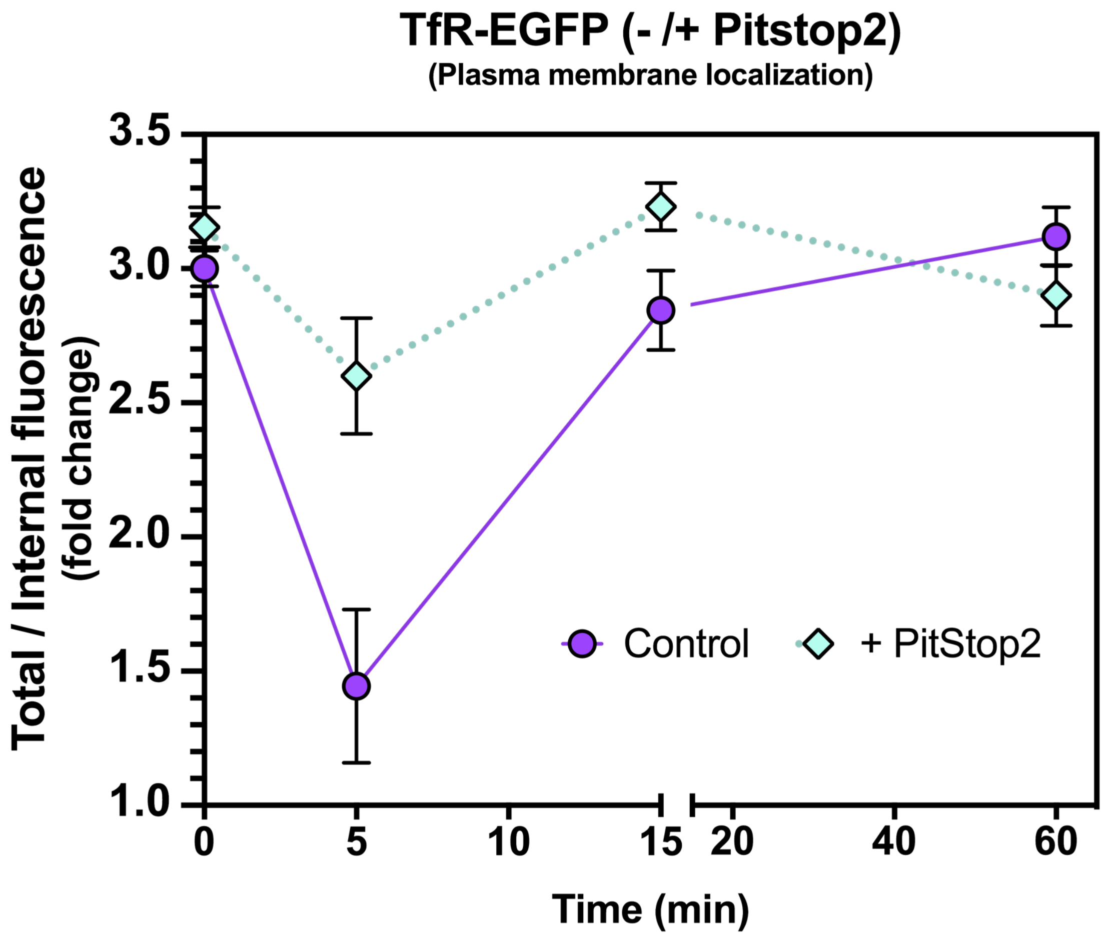

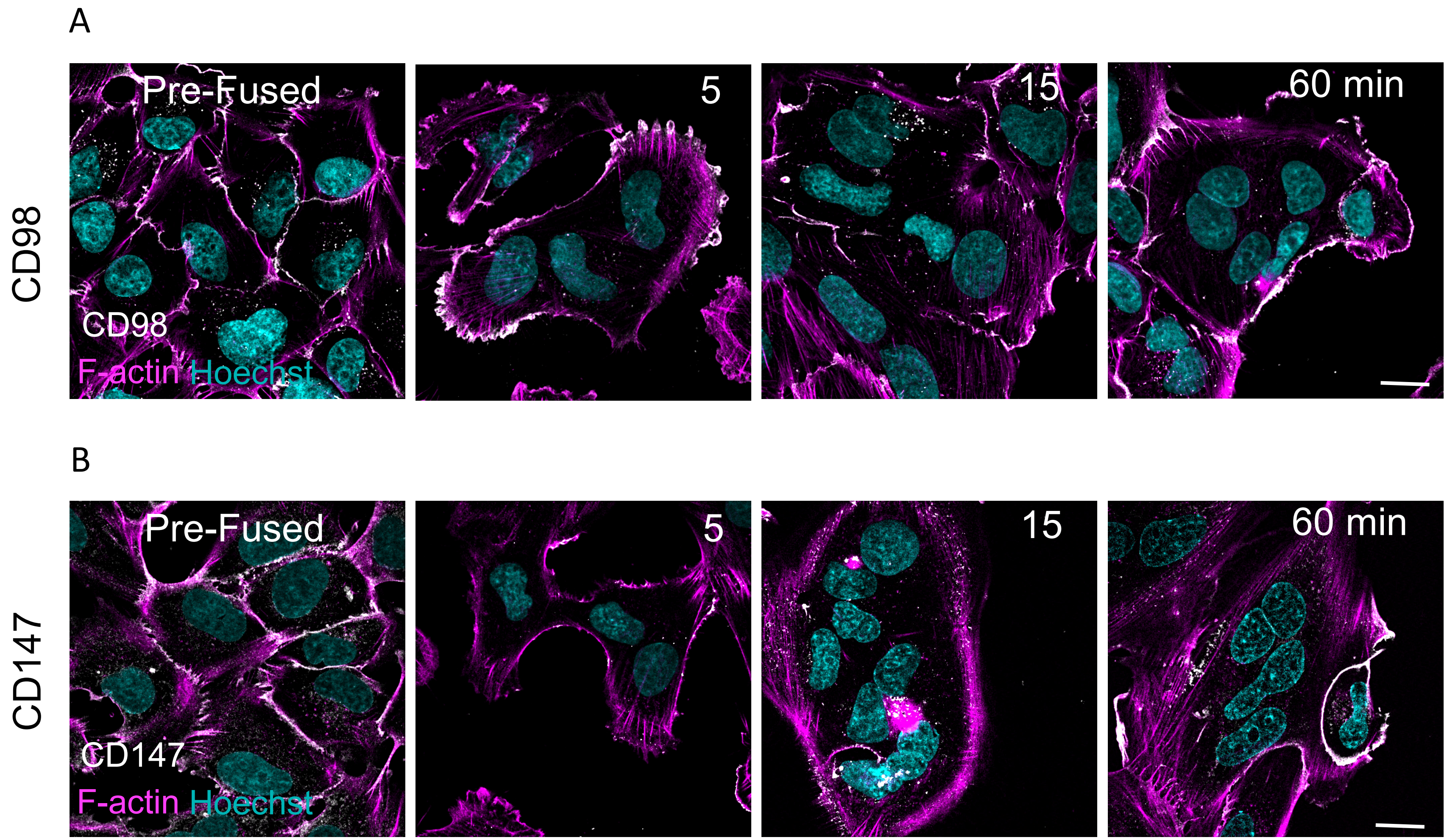
